# *APOE* Genotype and Biological Age Impact Inter-Omic Associations Related to Bioenergetics

**DOI:** 10.1101/2024.10.17.618322

**Authors:** Dylan Ellis, Kengo Watanabe, Tomasz Wilmanski, Michael S. Lustgarten, Andres V Ardisson Korat, Gwênlyn Glusman, Jennifer J. Hadlock, Oliver Fiehn, Paola Sebastiani, Nathan D. Price, Leroy Hood, Andrew T. Magis, Simon J. Evans, Lance Pflieger, Jennifer C. Lovejoy, Sean M. Gibbons, Cory C. Funk, Priyanka Baloni, Noa Rappaport

## Abstract

Apolipoprotein E (*APOE*) modifies human aging; specifically, the ε2 and ε4 alleles are among the strongest genetic predictors of longevity and Alzheimer’s disease (AD) risk, respectively. However, detailed mechanisms for their influence on aging remain unclear. Herein, we analyzed inter-omic, context-dependent association patterns across *APOE* genotypes, sex, and health axes in 2,229 community-dwelling individuals to test *APOE* genotypes for variation in metabolites and metabolite-associations tied to a previously-validated metric of biological aging (BA) based on blood biomarkers. Our analysis, supported by validation in an independent cohort, identified top *APOE*-associated plasma metabolites as diacylglycerols, which were increased in ε2-carriers and trended higher in ε4-carriers compared to ε3-homozygotes, despite the known opposing aging effects of the allele variants. ‘Omics association patterns of ε2-carriers and increased biological age were also counter-intuitively similar, displaying increased associations between insulin resistance markers and energy-generating pathway metabolites. These results provide an atlas of *APOE*-related ‘omic associations and support the involvement of bioenergetic pathways in mediating the impact of *APOE* on aging.

## Introduction

Aging is accompanied by a progressive decrease in physiological integrity, which results from the accumulation of damage in different molecular systems and is characterized by genomic instability, deregulated nutrient-sensing, mitochondrial dysfunction, and cellular senescence.^1^ Older age increases the risk of death and is the biggest predictor of neurodegenerative disorders such as Alzheimer’s disease (AD), which is the leading cause of dementia.^2^ We and others have shown that aging does not occur at the same rate for each individual, implying that a person’s chronological age (CA) is an imprecise measure of their biological age (BA).^3–5^ The difference (i.e., delta) between BA and CA (BA − CA) can be used to represent an individual’s health state normalized for their CA, with a negative delta signifying better health and a positive delta signifying worse health. BA and delta age are modifiable through lifestyle choices,^3^ and are of interest for designing, proposing, and evaluating wellness interventions.

The haplotypes of the human apolipoprotein E gene (APOE) exert strong, divergent effects on aging, with the ε4 allele being the greatest genetic predictor of late onset AD incidence,^6–8^ whereas the ε2 allele is protective against AD risk,^9,10^ and is a predictor of longevity independent of AD.^11,12^ However, despite APOE’s long established connection to AD incidence and longevity, the mechanisms underlying its apparent influence on aging and neurodegeneration remain largely uncharacterized. Recent research trends have supported metabolic and immuno-metabolic hypotheses of AD etiology, pointing to perturbations within mitochondrial function, impairments in glucose metabolism and other bioenergetic alterations both peripherally and within the brain as potentially causal mechanisms for dementia and the associated hallmark accumulation of amyloid beta (Aβ).^13–20^ We and others hypothesize that APOE could be partly responsible for the complex, interwoven shifts seen in aging and AD, with APOE ε4 influencing both brain and blood metabolomes.^16,21–23^ A recent case report of an individual homozygous for the APOE Christchurch mutation, resistant to a familial PSEN1 mutation, suggests APOE’s effects are upstream of amyloid production.^24^ This aligns with our recent analysis of the Alzheimer’s Disease Neuroimaging Initiative (ADNI) data showing that APOE’s reduced ability to off-load excess cholesterol, as well as the redistribution of cholesterol and other fatty acids across cell types in the brain, disrupts metabolic support for neurons by interfering with GPCR signaling in the astrocyte-neuron lactate shuttle.^25^ Understanding how different forms of APOE affect health throughout life and the context-dependency of its systemic influences on metabolism and aging could provide targets and help in preventing AD. Given the complexity of multifaceted phenotypes such as aging and AD, it is important to investigate beyond changes in individual measurements, and examine how networks of interacting biological features are altered. We thus set out to further understand the effects of APOE on system dynamics.

We studied multi-omic data from an AD-undiagnosed cohort of 2,229 community dwelling individuals aged 19-83, investigating the impact of APOE genotype and delta age on inter-omic associations (those spanning different types of molecular phenotypic data, for example between clinical chemistries and the metabolome). Our results indicate that APOE ε2 carriers and ε4 carriers display a similar increased abundance of plasma diacylglycerols (DAGs) and modified associations in bioenergetic pathways, including changes in ε2-carrying males resembling those of biologically older males. Our results provide an atlas for intervention targets to potentially reduce AD risk and promote longevity, and further contextualize the complex relationship between APOE, biological aging, and insulin resistance.

## Results

### Study design and cohort summary

This study aimed to analyze how APOE genotype and delta age (BA − CA) are associated with shifts in blood metabolomes and inter-omic associations in community-dwelling individuals without an AD diagnosis, using data from the Arivale^26^ and TwinsUK^27^ cohorts. (**Figure 1**).

**Figure 1:**
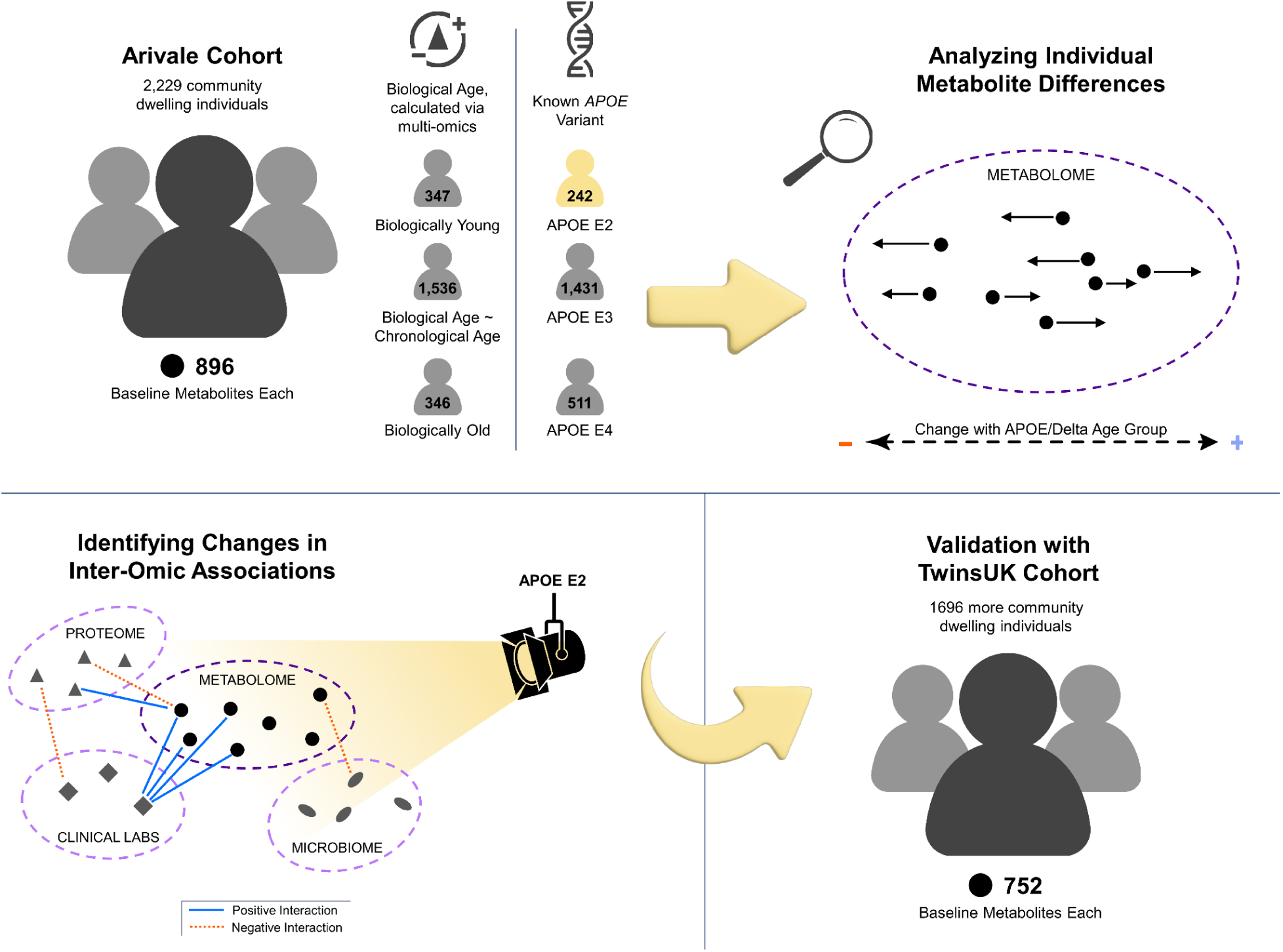
Study design to identify APOE genotype- and delta age-related alterations in the metabolome and inter-omic associations. Community dwelling individuals from the Arivale cohort were sorted based on delta age and APOE ε2 or ε4 carrier status. Metabolomic changes across APOE and delta age statuses were then analyzed. Finally, an inter-omic interaction analysis was performed to identify the effect modification of APOE or delta age status on inter-omic associations. These findings elucidate potential context-dependent relationships within APOE status and delta age group. Analyses were then repeated for validation with the TwinsUK cohort.

Differential metabolite abundances were first analyzed across APOE genotypes and delta age groups. This analysis was followed by an inter-omic interaction analysis, wherein the influence of a condition of interest, such as APOE status (APOE E2 for APOE ε2/ε2 or ε2/ε3; APOE E3 for ε3/ε3; or APOE E4 for ε3/ε4 and ε4/ε4) or delta age status, on the association between two analytes of different ‘omes was evaluated. The significant inter-omic interactions observed with each sex-stratified APOE status and delta age status were compared.

Baseline data from the Arivale Scientific Wellness dataset^26^ was used as a discovery cohort. The Arivale cohort, a former consumer-facing wellness company, consists of subscribers who were deeply-phenotyped and provided with personalized health coaching. Participants ranged from 19 to 83 (mean 46.6) years of age and represented the health of the communities they were drawn from^26^ (**Figure S1**). Multi-omic BAs were previously calculated,^3^ and delta age statuses were defined as biologically older for those with BA at least 7.5 years (∼one standard deviation) older than CA, and biologically younger for those with BA at least 7.5 years younger than CA for males and females. We did not observe differences in delta age across APOE status (**Figure S2**). We used data from TwinsUK as a validation cohort, which was originally intended to investigate rheumatologic diseases in identical twins in the United Kingdom, and has since expanded to encompass over 15,000 volunteer identical and non-identical twins.^27^ The subsection of the cohort with plasma metabolomics^28^ and clinical lab data available for use in this study was 96.4% female, 99.9% non-Hispanic White, and had an older population than Arivale with ages ranging 32 to 87 (mean 58.1) years at baseline. **Table 1** provides a demographic summary of the Arivale and TwinsUK cohorts. Using the same method as previously performed on Arivale data, a metabolomics-based BA was calculated for TwinsUK individuals in this study (Pearson’s r = 0.778 for females, 0.776 for males, see Methods and **Figure S3**), with the delta age status cutoff defined as 7.5 years for females and 5.0 years for males, reflective of their standard deviations. In TwinsUK, delta age was found to be increased in APOE E4 in females but decreased in APOE E4 in males (**Figure S4**).

**Table 1:** Summary of Arivale and TwinsUK cohorts. Continuous data (age, BMI, delta age) are reported as ‘mean (range)’. All other fields are reported as percentage, with E2 representing ε2/ε2 and ε2/ε3, E3 representing ε3/ε3, and E4 representing ε3/ε4 and ε4/ε4. Data reported for each cohort is for individuals directly prior to use for interaction analysis. Baseline data was used for both Arivale and TwinsUK. Delta age data of the Arivale cohort was derived from the combined BA model predictions.

|  | Arivale |  | TwinsUK |  |
| --- | --- | --- | --- | --- |
|  | Females | Males | Females | Males |
| **N** | 1403 | 826 | 1635 | 61 |
| **Age (years)** | 46.6 (21.0-83.0) | 44.5 (19.0-80.0) | 51.4 (32.9-73.7) | 51.1 (33.6-58.5) |
| **Race/Ethnicity** |  |  |  |  |
| **(% non-Hispanic Whites)** | 74.4% | 66.0% | 99.9% | 98.4% |
| **BMI (kg/m<sup>2</sup>)** | 28.6 (16.9-62.3) | 27.7 (18.0-61.5) | 25.4 (16.5-48.2) | 26.6 (20.3-36.3) |
| **% Cholesterol Medication Use** | 9.1% | 14.0% | 3.2% | 3.3% |
| **% APOE E2** | 11.3% | 10.2% | 13.1% | 9.8% |
| **% APOE E3** | 63.6% | 65.3% | 63.1% | 52.5% |
| **% APOE E4** | 22.7% | 23.2% | 21.3% | 37.7% |
| **Delta Age (years)** | 0.5 (-28.0-+41.2) | -0.4 (-35.0-43.1) | -2.6 (-25.4-+22.4) | 1.8 (-8.3-+16.7) |

### Diacylglycerols and plasmalogens are the metabolites most significantly associated with APOE genotype and delta age status

To analyze the associations of APOE and delta age group with the metabolome, we constructed two sets of generalized linear models (GLMs) for each metabolite: one using metabolite abundance as the dependent variable, APOE E2 and E4 statuses as the independent variables, and covariates (age, BMI, use of cholesterol medications, sex, and first two genetics principal components) and the other using biologically younger and older delta age groups in place of APOE status (see Methods). **Figure 2** depicts the distribution of *β*-coefficient estimates and their significance for the experimental variables from these models. Out of 896 metabolites, 87 differential metabolite abundance GLMs had APOE E2 *β*-coefficient estimates with pre-adjusted *p* < 0.05, while 67 had pre-adjusted *p* < 0.05 for E4. After adjusting for false-discovery rate (FDR, Benjamini-Hochberg method with 5% FDR used throughout), 20 metabolites retained significant associations at pFDR < 0.1 for APOE E2. Top positively APOE E2-associated metabolites included DAGs such as linoleoyl-arachidonoyl-glycerol (18:2/20:4) [1]*

**Figure 2:**
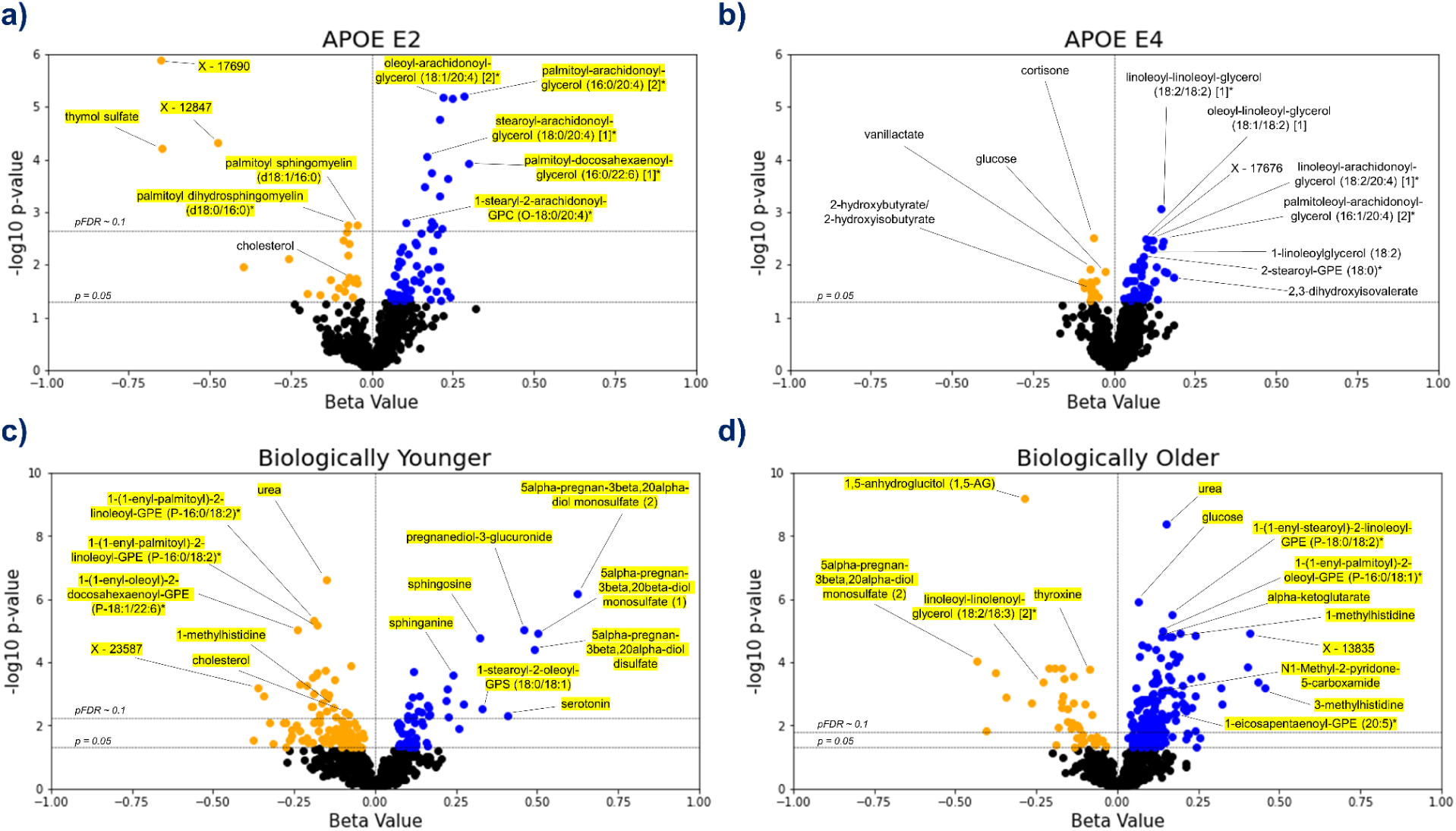
Lipids are the main APOE- and delta age-associated metabolites. (**a**–**d**) Volcano plots for the APOE E2 (**a**), APOE E4 (**b**), biologically younger (**c**), or biologically older (**d**) groups. For each metabolite, presented are the *β*-coefficient estimate and its log10 *p*-value from the GLM including metabolite abundance as the dependent variable, group statuses as the independent variables, and age, BMI, use of cholesterol medications, sex, and first two genetics principal components as the covariates (see Methods). Blue data points indicate a positive association between metabolite and test group with pre-adjusted *p* < 0.05, whereas orange points indicate a negative pre-adjusted association. Yellow highlighting indicates significance after multiple hypothesis testing (pFDR < 0.1, Benjamini–Hochberg method). *n* = 896 metabolites.

(*β*-coefficient estimate = 0.248), palmitoyl-arachidonoyl-glycerol (16:0/20:4) [2]* (*β* = 0.285), and oleoyl-arachidonoyl-glycerol (18:1/20:4) [2]* (*β* = 0.219) (all pFDR = 1.58e-3), and top negatively APOE E2-associated metabolites included sphingolipids such as palmitoyl dihydrosphingomyelin (d18:0/16:0)* (*β* = −0.074) and palmitoyl sphingomyelin (d18:1/16:0) (*β* = −0.044) (both pFDR = 0.089). Thymol sulfate was another top negatively E2-associated metabolite (*β* = −0.647, pFDR = 7.72e-3). For APOE E4, DAGs trended toward positive associations though were not significant after FDR-adjustment, including linoleoyl-linoleoyl-glycerol (18:2/18:2) [1]* (*β* = 0.144, pre-adjusted *p* = 8.58e-4), oleoyl-linoleoyl-glycerol (18:1/18:2) [1] (*β* = 0.100, *p* = 3.22e-3) and linoleoyl-arachidonoyl-glycerol (18:2/20:4) [1]* (*β* = 0.120, *p* = 3.45e-3) (all had pFDR = 0.514). Of the 13 DAGs included in the Arivale dataset, all were positively associated pre-adjustment (*p* < 0.05) with E2 (8 out of 13 also with pFDR < 0.1, post-adjustment), while 4 were positively associated pre-adjustment with E4. Those DAGs associated with E4 pre-adjustment mostly contained linoleoyl acyl groups, while those with stronger significant associations with E2 tended to contain more palmitoyl, oleoyl, and stearoyl groups. We further performed enrichment analysis for the metabolites significantly (pFDR < 0.1) positively or negatively associated with APOE E2 and E4 using the sub-pathways annotated by the Metabolon platform (**Supplementary Table 1**). DAGs were enriched in the positive APOE-associated metabolites for the E2 group (pFDR = 2.55e-12). We also performed enrichment analysis on those metabolites with pre-adjusted (*p* < 0.05) positive and negative associations with APOE E2 and E4 (**Supplementary Table 2**). In addition to the enrichment of DAGs, these sets of associations with pre=adjusted *p* < 0.05 were enriched for plasmalogens and long chain fatty acids in the positive associations with APOE E2 (pFDR = 0.075 for both). The sphingolipid metabolism sub-pathway was enriched in negative APOE E2 associations (pFDR = 2.33e-5). For the positive associations with APOE E4, lysolipids were enriched (pFDR = 0.091). DAGs were also narrowly outside FDR significance (pFDR = 0.111, *p* = 2.61e-3) for positive associations in APOE E4, showing a similar trend to APOE E2 despite the expected opposite effect of these genotypes.

For the delta age analyses using GLMs (**Figure 2c, d**), 51 metabolites for the biologically young group were significant at pFDR < 0.1 (158 metabolites had pre-adjusted *p* < 0.05), while 143 were significantly associated with the biologically old group (227 were associated pre-adjustment). Urea was the top metabolite negatively associated with the biologically young group (*β* = −0.150, pFDR = 2.22e-4) and the second top metabolite positively associated with the biologically old group (*β* = 0.151, pFDR = 1.80e-6). For the biologically young, we observed positive associations for steroid metabolites such as 5alpha-pregnan-3beta,20alpha-diol monosulfate (2) (*β* = 0.624, pFDR = 2.89e-4) and pregnanediol-3-glucuronide (*β* = 0.459, pFDR = 1.38e-3) among the most significant, as well as sphingosine (*β* = 0.322, pFDR = 1.86e-3) and sphinganine (*β* = 0.240, pFDR = 0.016). For the biologically old, 1,5-anhydroglucitol (1,5-AG), a known marker for glycemic control that is inversely associated with diabetes risk,^29,30^ had the most significant *β*-coefficient estimate and was negatively associated (*β* = −0.287, pFDR = 5.64e-7). Glucose (*β* = 0.065, pFDR = 3.65e-4) and alpha-ketoglutarate (*β* = 0.135, pFDR = 1.30e-3), central bioenergetic metabolites, were the third and tenth most significant metabolites for the biologically old, both positively associated, while neither had FDR significant associations for the biologically young. 1-methylhistidine was also near the top of the list of significance for both groups, being negatively associated with the biologically young (*β* = −0.131, pFDR = 0.068) but positively associated with the biologically old (*β* = 0.167, pFDR = 1.30e-3). The following enrichment analysis on the significant (pFDR < 0.1) associations with delta age statuses (**Supplementary Table 1**) revealed that plasmalogens, phospholipids crucial for cell membrane integrity and linked to important roles in cognitive health and neurological function, were the most significantly enriched sub-pathway in metabolites negatively associated with the biologically young (pFDR = 4.28e-5) and in metabolites positively associated with the biologically old (pFDR = 8.52e-8). Steroid metabolites were also enriched in positive associations with the biologically young (pFDR = 0.088), and the polyamine metabolism subpath was enriched in the negative associations with the biologically old (pFDR = 0.030).

Between the biologically young and old, 21 metabolites were FDR-significant for both groups, all having diverging associations. Comparing the analyses of APOE and delta age status, two of the metabolites significantly (pFDR < 0.1) associated with APOE E2 were also significantly (pFDR < 0.1) associated with the biological old status: linoleoyl-linoleoyl-glycerol (18:2/18:2) [1]* was positively associated with APOE E2 but negatively associated with the biological old, while palmitoyl dihydrosphingomyelin (d18:0/16:0)* was negatively associated with APOE E2 but positively associated with the biological old. Several metabolites have associations with pre-adjusted *p* < 0.05 overlapping across experimental groups: 13 metabolites are associated concordantly with APOE E2 and biologically younger individuals, while 4 metabolites are associated discordantly. Between APOE E2 and biologically older individuals, 12 metabolites have concordant associations while 8 are discordant. For APOE E4 and biologically younger individuals, 1 metabolite is concordantly associated while 5 are discordantly associated. Finally, between APOE E4 and biologically older individuals, 14 metabolites have concordant associations and 7 have discordant associations. In a sex-stratified analysis, biologically old males and females also shared a significant (pFDR < 0.1) negative association with 1,5-AG and significant (pFDR < 0.1) positive associations with urea, glucose, and 1-palmitoyl-2-oleoyl-GPC (16:0/18:1) (**Supplementary File 1**).

### Bioenergetic analyte associations are modified by APOE and delta age status

To explore systemic and context-dependent omics changes associated with APOE and delta age, we assessed the associations between 509,360 inter-omic pairs (being of different ‘omes) of analytes across the plasma metabolome, plasma proteome, gut microbiome, and clinical chemistries using an analyte-by-experimental group (i.e., APOE E2, APOE E4, biologically young, or biologically old) interaction term in the GLMs for each pair (see Methods). This type of statistical test, called an interaction analysis, assesses whether the relationship between two analytes is dependent on a third variable (in this case, the experimental group). The association between two analytes can be positively or negatively modified by a third variable, such as the association between glucose and *Klebsiella* being more positive in APOE E2 than in E3 in males in this study. Other recent works have successfully employed interaction analyses to identify multi-omic differences in COVID-19 disease states^31^ and to examine how the associations between proteins and AD incidence are modified by APOE ε4 carriership,^32^ as examples.

The ten analyte pairs with the lowest *p*-values for the interaction term in each sex-stratified experimental group model are presented in **Table 2**, and the top ten pairs for models using allele dosage and continuous delta age are presented in **Supplementary Table 3**. Individual associations, or lack thereof, between analytes and either APOE or delta age did not appear to influence whether analytes were identified in the interaction analysis, suggesting that the interaction analysis provides a unique layer of information. Indeed, out of the 79 metabolites appearing in the top twenty significant analyte pairs of all the APOE-related interaction analyses, 65 were detected as significant exclusively in the interaction analysis and not (*p* > 0.05) in the earlier differential metabolite abundance analysis, including fumarate, malate, and ribitol.

**Table 2:**
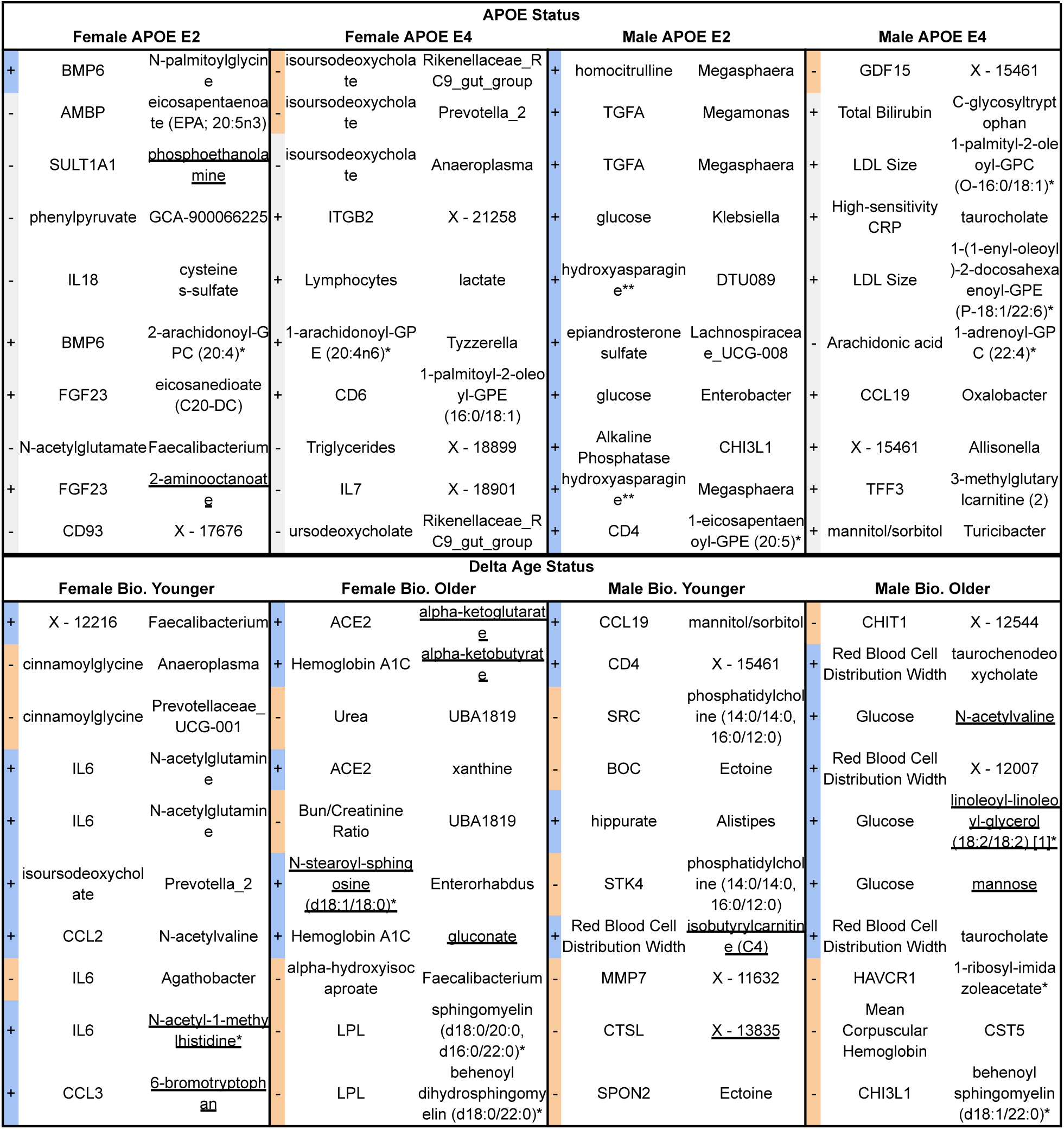
Top ten inter-omic analyte pair associations modified by each APOE and delta age status, stratified by sex. For each test group, the ten analyte pairs with the lowest *p*-values for the interaction term representing the modification of APOE or delta age status on the association between the two analytes are tabulated. Blue and orange highlighting indicate significant positive and negative interaction terms, respectively, while gray indicates pFDR > 0.1 (**Supplementary File 2** for full data). Underlining indicates a metabolite associated with the experimental group in the analysis of differential metabolite abundance (with pre-adjusted *p* < 0.05).

Similarly, for the delta age-related interaction analyses, 37 out of the 56 metabolites in the top twenty pairs of each were exclusively significant in the interaction analyses, including pyruvate, lactate, and glutamate. For APOE E2 males, 60 significant (pFDR < 0.1) interactions were identified, including positively modified associations between glucose and *Klebsiella*; triglycerides and ribitol; as well as both glucose and phenol sulfate. APOE ε2 allele dosage significantly modified 17 inter-omic associations, including positively modified bioenergetic associations such as HbA1c with malate and fumarate as well as glucose with aconitate [cis or trans], as well as negatively modified associations between both LDL particle number and LDL small particle number with LDLR. APOE ε4 allele dosage significantly modified 5 inter-omic associations, including negatively modifying the association between isoursodeoxycholate and both *Rikenellaceae RC9 gut group* and *Prevotellaceae UCG-001*. Biologically younger males and females exhibited 28 and 16 significant interaction pairs, respectively, with SRC (proto-oncogene non-receptor tyrosine kinase Src protein) appearing frequently in negatively modified associations with several metabolites in the males and IL6 appearing in positively modified associations with N-acetylglutamine but negatively modified associations with *Agathobacter*. Biologically older males and females significantly modified 526 and 89 associations, respectively, including many positively modified associations containing glucose or HbA1c from the clinical chemistries with bioenergetic metabolites such as pyruvate and alpha-ketoglutarate. Continuous delta age showed a similar signature in its 840 significantly modified associations, including positively modifying the association between HbA1c and pyruvate, mannose, lactate, CD163, gluconate, and fructose.

### Comparison of inter-omic signatures of APOE and delta age status

Analyte pairs having interaction terms with pFDR < 0.1 for each subgroup for both sets of interaction analyses were compared to identify similarities and differences between the contextual manifestation of inter-omic associations of APOE and delta age groups in both males and females. We directly compared the inter-omic association signatures between these groups to characterize the perturbations in these complex systems and to capture the essence of age-related metabolic shifts by pinpointing the changes in associations that converge or diverge across conditions. **Figure 3** highlights key modified inter-omic associations and compares the association signatures across APOE E2 males and biologically older males as well as across biologically older males and females. APOE E2 males and biologically older males shared four significant interactions, all being positively modified in both groups: those between hydroxyasparagine** and *Megasphaera*; FST (follistatin protein) and laureate (12:0); HbA1c and phenol sulfate; and glucose and phenol sulfate. Biologically older males and females shared five pairs, with all the associations being positively modified in both: HbA1c and pyruvate; HbA1c and mannose; glucose and HGF (hepatocyte growth factor); glucose and CD163; and glucose and X - 16087 (unknown metabolite). In addition to these modified associations directly shared between biologically old males and females, each group had associations with metabolites from similar pathways modified. For instance, biologically older males showed positively modified associations between HbA1c and aspartate, glutamate, lactate, mannose, and laureate (12:0), while biologically older females had positively modified associations between HbA1c and alpha-ketoglutarate, margarate (17:0), and taurine. The only other significantly modified association overlapping in multiple of these groups was that between isoursodeoxycholate and *Prevotella 2*, which was positively modified in biologically younger females but negatively modified in APOE E4 females. Finally, for the models testing APOE allele dosage and continuous delta age, both APOE ε2 allele dosage and delta age positively modified the associations between glucose and aconitate [cis or trans], and glucose and 3-hydroxy-2-ethylpropionate (**Figure S5**). More comprehensive results of the inter-omic interaction analyses are included in **Supplementary File 2**.

**Figure 3:**
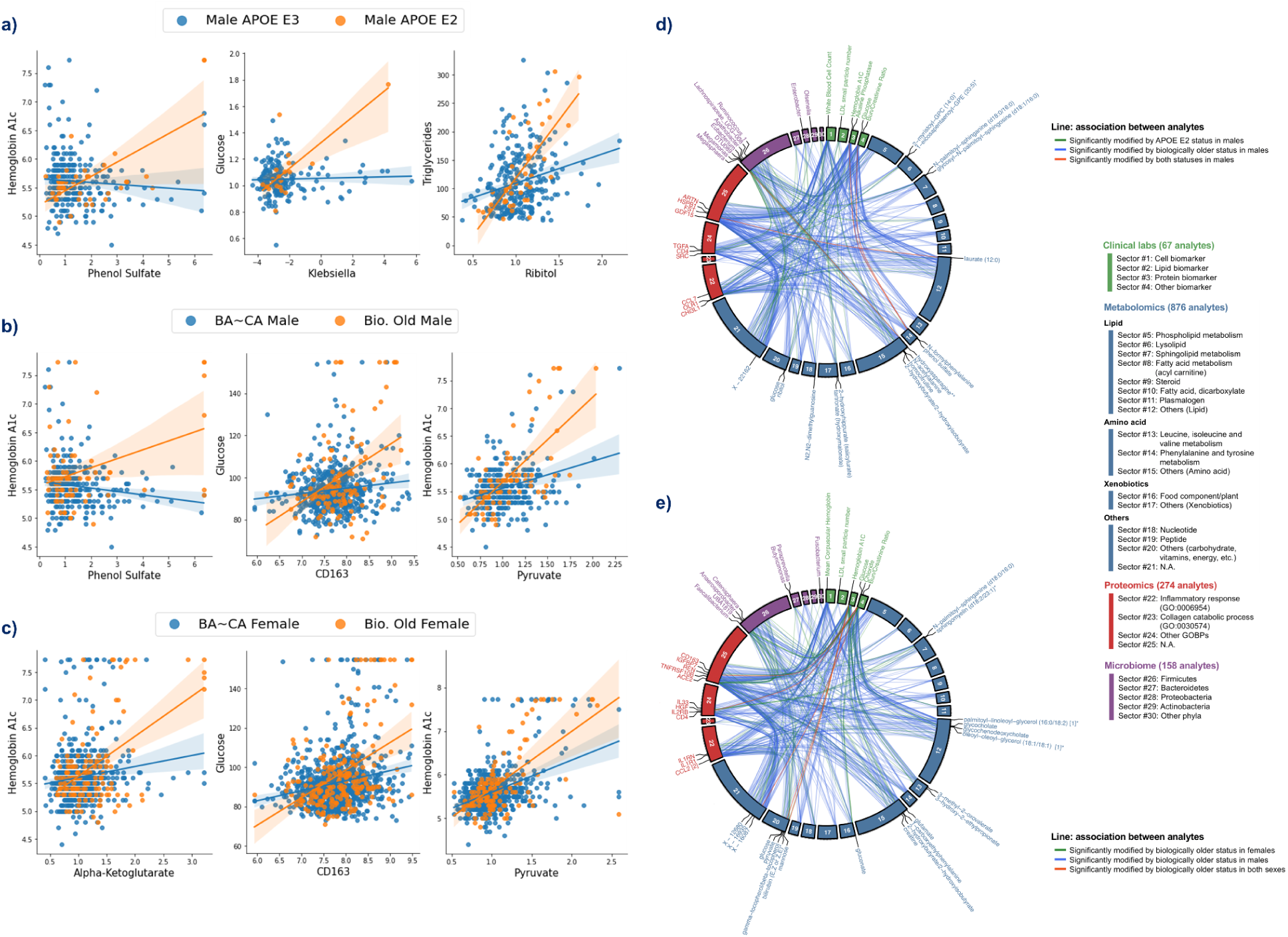
Biologically older males show similar multi-omic association signatures to APOE E2 males and biologically older females, particularly within central bioenergetic analytes. (**a-c**) Scatter plots of inter-omic analyte pairs with associations significantly modified by APOE E2 in males (**a**), and by biological oldness in males (**b**) and females (**c**). Line indicates simple linear regression, with shading indicating the 95% confidence interval. (**d-e**) Circos plots depicting the shared analyte associations (pFDR < 0.1, Benjamini-Hochberg method) between male APOE E2 and biologically older males (**d**), and between biologically older males and females (**e**). Associations specific to one group are connected with green and blue lines, whereas significant concordant associations shared in both groups are presented in red lines. Analyte nodes in associations significant to both groups are labeled.

### Validation analyses in TwinsUK

In the individual GLMs, out of 752 total metabolites (with 547 overlapping those in Arivale), 74 associations with APOE E2 and 80 with APOE E4 had *p* < 0.05. After FDR adjustment, three metabolites had pFDR < 0.1 for APOE E2: cholesterol (*β* = −0.089) and sphingomyelin (d18:1/20:0, d16:1/22:0)* (*β* = −0.086) were decreased, and an unidentified metabolite was increased (*β* = 0.213) (all pFDR = 0.059) (**Figure 4**). Consistent with the Arivale finding, lipids appeared amongst the most significant APOE-associated metabolites, and DAGs were significantly enriched in the metabolites with *p* < 0.05 for associations with APOE E2 and E4 (pFDR = 1.10e-3 and 0.036, respectively) (**Supplementary Table 2**). Analytes involved in sphingolipid metabolism were also significantly enriched in the metabolites with *p* < 0.05 negatively associated with APOE E2 (pFDR = 2.95e-4), as in Arivale (pFDR = 2.33e-4).

**Figure 4:**
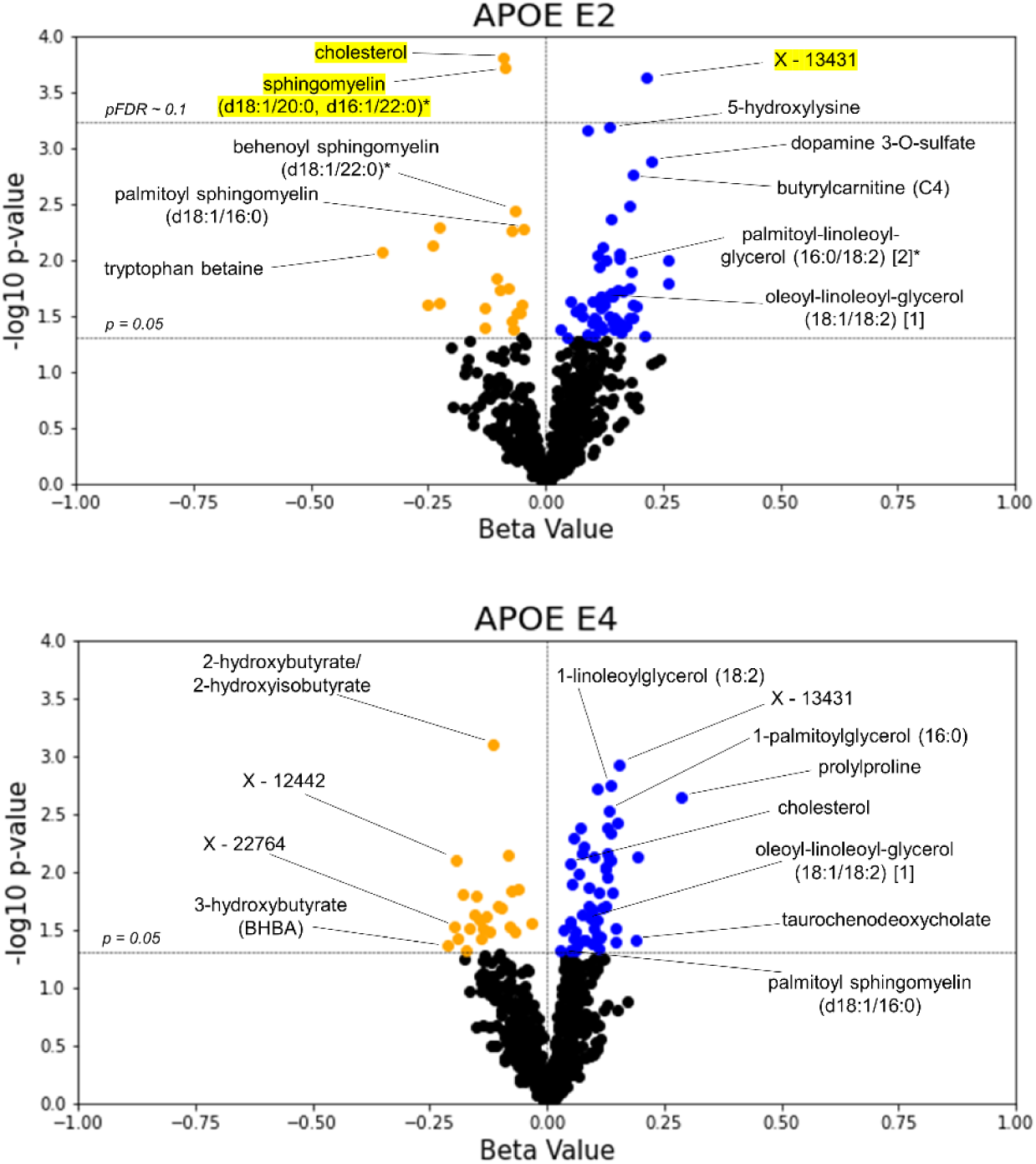
TwinsUK validates lipids as top APOE associated metabolites. The *β*-coefficient estimates for the APOE E2 and E4 groups are plotted against their -log10 pre-adjusted *p*-value from the metabolite GLMs. Blue data points indicate a positive association between metabolite and test group with pre-adjusted *p* < 0.05, whereas orange points indicate a negative pre-adjusted association. Yellow highlighting indicates significance after multiple hypothesis testing (pFDR < 0.1, Benjamini-Hochberg method).

We further explored direct comparisons of the metabolites with pre-adjusted *p* < 0.05 in both the Arivale and TwinsUK cohorts (**Table 3**). Out of the 547 overlapping metabolites tested in Arivale and TwinsUK, nine metabolites had *p* < 0.05 in both cohorts with the same direction for APOE E2: four DAGs, cholesterol, two sphingomyelins, and two other lipids. One metabolite, 3-methylcatechol sulfate, had *p* < 0.05 in both cohorts but in the opposite direction for APOE E2. For APOE E4, 11 metabolites had *p* < 0.05 in both cohorts with the same directional difference, including three DAGs, two monoacylglycerols, and 2-hydroxybutyrate/2-hydroxyisobutyrate. The metabolite associations with delta age groups were not tested in the TwinsUK cohort, because the BA model was able to be generated only from the metabolomics data.

**Table 3:** Comparison of significant metabolites in differential abundance GLM tests for APOE across Arivale and TwinsUK. Metabolites with pre-adjusted *p* < 0.05 for at least two out of four APOE beta values in metabolite abundance GLMs across tests for Arivale and TwinsUK are reported. ‘+’ indicates a positive beta value, ‘-’ indicates a negative beta value, and ‘ns’ indicates nonsignificance. Metabolites missing from one of the two datasets are denoted. Metabolites are sorted alphabetically. Yellow highlighting denotes pFDR < 0.1.

| Metabolite | Arivale E2 | Arivale E4 | TwinsUK E2 | TwinsUK E4 |
| --- | --- | --- | --- | --- |
| 1-(1-enyl-stearoyl)-2-arachidonoyl-GPC (P-18:0/20:4) | + | - | ns | ns |
| 1-(1-enyl-stearoyl)-2-dihomo-linolenoyl-GPE (P-18:0/20:3)* | + | - | Not in Dataset | Not in Dataset |
| 1-(1-enyl-stearoyl)-2-oleoyl-GPE (P-18:0/18:1) | + | ns | + | ns |
| 1-docosapentaenoyl-GPC (22:5n3)* | ns | + | ns | + |
| 1-linoleoylglycerol (18:2) | + | + | ns | + |
| 1-oleoylglycerol (18:1) | + | + | ns | + |
| 1-stearoyl-2-arachidonoyl-GPE (18:0/20:4) | + | ns | + | ns |
| 1-stearoyl-2-linoleoyl-GPE (18:0/18:2)* | ns | + | ns | + |
| 1-stearoyl-GPE (18:0) | ns | + | ns | + |
| 1-stearyl-2-arachidonoyl-GPC (O-18:0/20:4)* | + | - | Not in Dataset | Not in Dataset |
| 2-hydroxybutyrate/2-hydroxyisobutyrate | ns | - | ns | - |
| 2-linoleoyl-GPE (18:2)* | ns | + | ns | + |
| 2-stearoyl-GPE (18:0)* | ns | + | ns | + |
| 3-(3-hydroxyphenyl)propionate | Not in Dataset | Not in Dataset | - | - |
| 4-methylcatechol sulfate | - | ns | + | ns |
| cholesterol | - | ns | - | + |
| diacylglycerol (16:1/18:2 [2], 16:0/18:3 [1])* | + | + | Not in Dataset | Not in Dataset |
| isoleucylvaline | Not in Dataset | Not in Dataset | + | + |
| linoleoyl-arachidonoyl-glycerol (18:2/20:4) [1]* | + | + | Not in Dataset | Not in Dataset |
| linoleoyl-linoleoyl-glycerol (18:2/18:2) [1]* | + | + | Not in Dataset | Not in Dataset |
| oleoyl-arachidonoyl-glycerol (18:1/20:4) [1]* | + | + | Not in Dataset | Not in Dataset |
| oleoyl-arachidonoyl-glycerol (18:1/20:4) [2]* | + | + | Not in Dataset | Not in Dataset |
| oleoyl-linoleoyl-glycerol (18:1/18:2) [1] | + | + | + | + |
| oleoyl-linoleoyl-glycerol (18:1/18:2) [2] | + | + | + | + |
| palmitoleoyl-arachidonoyl-glycerol (16:1/20:4) [2]* | + | + | Not in Dataset | Not in Dataset |
| palmitoleoyl-linoleoyl-glycerol (16:1/18:2) [1]* | + | + | Not in Dataset | Not in Dataset |
| palmitoyl dihydrosphingomyelin (d18:0/16:0)* | - | ns | - | ns |
| palmitoyl sphingomyelin (d18:1/16:0) | - | ns | - | + |
| palmitoyl-arachidonoyl-glycerol (16:0/20:4) [2]* | + | + | Not in Dataset | Not in Dataset |
| palmitoyl-linoleoyl-glycerol (16:0/18:2) [1]* | + | + | + | + |
| palmitoyl-linoleoyl-glycerol (16:0/18:2) [2]* | + | ns | + | ns |
| pyrraline | Not in Dataset | Not in Dataset | + | + |
| X - 11491 | Not in Dataset | Not in Dataset | + | + |
| X - 11795 | ns | + | ns | - |
| X - 13431 | ns | ns | + | + |
| X - 24065 | Not in Dataset | Not in Dataset | - | + |

In the inter-omic interaction analysis for metabolomics and clinical lab measures, those pairs significant in Arivale and measured in the TwinsUK were tested along with 965 additional pairs (5 chemistries by 193 metabolites) containing analyte groups frequently appearing in Arivale hits across experimental groups and related to bioenergetics and lipid metabolism (see **Methods**).

Biologically young males showed the largest number of significant (pFDR < 0.1) hits, with 48 negatively modified associations mainly between phospholipids and LDL or total cholesterol and 3 positively modified associations, though low numbers of biologically younger males seemed to drive these associations. Biologically older males showed a negatively modified association between LDL and palmitoyl-linoleoyl-glycerol (16:0/18:2) [2]*. Biologically older females showed a negative modification on the association between glucose and isovalerylcarnitine (C5). APOE E2 positively modified the association between HDL and trimethylamine N-oxide but negatively modified the association between LDL and 1-palmitoyl-2-stearoyl-GPC (16:0/18:0) in females. The APOE ε2 allele, regardless of sex, significantly modified these same associations as well as negatively modified the associations between 1,2-dipalmitoyl-GPC (16:0/16:0) and both LDL and total cholesterol. Finally, in females APOE E4 positively modified the associations between isovalerylglycine and both total cholesterol and LDL; negatively modified the association between total cholesterol and sphingomyelin (d18:1/20:1, d18:2/20:0)*; and negatively modified the associations between blood glucose and both leucine and isoleucine. Representative modified associations are highlighted in **Figure S6** while full results are provided in **Supplementary File 4**.

With the differences in composition between the cohorts, no significant interactions in Arivale were able to be validated with FDR-significance. However there were a few interactions in TwinsUK having pre-adjusted *p* < 0.05 that corroborated with observed significant interactions in Arivale. The associations between glucose and choline, valine, aspartate, leucine, glutamate, and palmitoyl-linoleoyl-glycerol (16:0/18:2) [2]* were observed to be positively modified by biological oldness in males for both cohorts, all with pre-adjusted *p* < 0.02 (max pFDR of 0.37) in TwinsUK. In females, biological oldness positively modified the association between glucose and fructose with *p* = 0.015 (pFDR = 0.529) in TwinsUK. Increasing delta age in TwinsUK positively modified associations between triglycerides and both lactate (*p* = 3.00e-4, pFDR = 0.209) and glucose (*p* = 7.61e-3, pFDR = 0.474). Finally, APOE E2 positively modified the association between triglycerides and ribitol (*p* = 5.91e-3, pFDR = 0.349) in TwinsUK males.

## Discussion

In this study, we leveraged two deeply phenotyped wellness cohorts of 2,229 (Arivale) and 1,696 (TwinsUK) individuals to analyze the systemic interplay between APOE genotype, delta age, sex, the blood metabolome, clinical chemistries, the proteome, and the microbiome. Our main findings include: (1) a resemblance between APOE E2 and E4-associated changes in blood metabolomics with increased DAG abundance, consistent with prior studies^22,23^ and confirming APOE’s role in bioenergetics and potentially insulin resistance via altered lipid metabolism; (2) inter-omic associations in males and females are more similarly altered in a biologically older state than a biologically younger state, highlighting the importance of context-dependence; and (3) ‘omics associations between central bioenergetic analytes such as HbA1c, glucose, and glycolysis/TCA metabolites as well as lipids are similarly modified in APOE E2 and increased delta age, suggesting that APOE may systematically influence bioenergetic pathways, consistent with metabolic hypotheses of AD.

DAGs were among the most significant individual metabolites altered in APOE E2 and E4 in Arivale, and were positively associated in both groups as well as in the cohorts. DAGs have previously been shown to be increased in human plasma for ε2 carriers^23^ and are here observed to be increased in ε4 carriers (pre-FDR-adjustment). In contrast, DAGs have also been previously shown to be increased in ε2 carriers but decreased in ε4 carriers in the entorhinal cortex in mice models.^22^ This discrepancy of APOE E4’s potential influence on DAG levels between the blood and brain may point to differences in DAG transport and sequestering as well as metabolism across APOE, or potentially highlight the limitations of mouse models to accurately reflect humans. More likely, differences in the brain from the periphery may be due to the fact that the brain only has a single particle system for re-distributing and off-loading cholesterol (APOE), while the periphery has a two particle system: APOB for distributing cholesterol to cells, and APOA1, which helps transport excess cholesterol back to the liver.^33,34^ DAGs have also been observed to increase in both the plasma and the neocortex of AD patients, relative to controls.^35^ DAGs are a major hallmark of overall lipid oxidation, indicative of lipase acting on triglycerides. DAGs act as secondary messengers to activate protein kinase C (PKC) and thus propel cascades producing ROS and inflammatory cytokines,^36^ are associated with insulin resistance,^36,37^ and have been suggested as biomarkers for sustained immune activation.^35^ However, it is worth noting that different DAGs may have different effects based on the different isoforms and acyl groups present. A recent lipidomic study found overall plasma DAG levels to be positively correlated with higher steady-state plasma glucose levels, indicative of insulin resistance, yet also found DAGs to be negatively associated with age in participants with insulin resistance.^38^ High associations between plasma DAG levels and APOE isoforms suggests the lipid metabolism modulation by E2 (and potentially E4) increases plasma DAG accumulation, thereby potentially influencing insulin sensitivity and glucose uptake, even in a wellness state.

The similar pattern of elevated DAG species in both E2 and E4 is unexpected, given their typically opposing effects on aging in later decades of life. One reason may be that DAGs containing different acyl groups were variably associated with APOE E2 and E4. E2 was strongly associated with DAGs with palmitoyl and oleoyl residues. Palmitic and oleic acids are the most common saturated and monounsaturated fatty acids, respectively, and can both be synthesized *de novo* in humans or obtained via the diet, palmitic acid through meat and dairy or palm oils, and oleic acid largely from olive oil.^39,40^ Generally, increased palmitic acid is associated with poor health outcomes including inflammation, insulin resistance, and mitochondrial dysfunction,^39^ whereas oleic acid combats these effects and is associated with a healthier profile.^40^ More broadly, increased levels of circulating fatty acids related to *de novo* lipogenesis are associated with increased T2D incidence.^41^ E4s on the other hand tended to be more associated, albeit pre-FDR-adjustment, with those containing linoleoyl groups. Linoleic acid is the most commonly consumed polyunsaturated fatty acid, obtained exclusively in the diet, largely from vegetable oils.^42^ Some overlapping DAG species were however associated with both E2 and E4. This similarity might be due to the APOE isoforms distinct transport mechanisms, with E4 preferentially binding larger fat particles such as very low density lipoprotein (VLDL) and increased binding affinity to low density lipoprotein receptor, contrasting with E2.^43–45^ Both alleles might disrupt lipid transport or metabolism, leading to increased DAG as a common feature of inefficiency. Additionally, while E2 is generally seen as beneficial and E4 as harmful, their effects are complex and not strictly opposite. For instance, E2 is linked to certain vascular and cervical disorders, while E4 offers some protection against diseases like type 2 diabetes (T2D) and obesity,^46^ and these roles vary by sex and ancestry. Further research is needed to understand these mechanisms.

Our finding that 1-methylhistidine was significantly negatively associated with the biologically young but positively associated with the biologically old is interesting as a recent study identified the importance of histidine methylation in a subunit of mitochondrial complex I, NDUFB3, by METTL9 methyltransferase.^47^ Mitochondrial complex I activity and production of ROS has been studied in the context of longevity and neurodegenerative disorders including Parkinson’s and Alzheimer’s.^48–51^ Paradoxically, partial inhibition of mitochondrial complex I by the compound CP2 is beneficial for APP/PS1 mice that accumulate amyloid, restoring their cognitive function, as well as other markers of pathology, while treatment with CP2 in mice control animals shows no significant improvement.^52^ This is consistent with the directionality of our observation of 1-methylhistidine being associated with increasing delta age, thus suggesting decreased METTL9 activity producing 1-methylhistidine and activating mitochondrial complex I is beneficial and associated with a lower delta age.

Plasmalogens, a subclass of glycerophospholipids found in high amounts in the brain, heart, and myelin, were enriched in positive associations with E2 and biologically older individuals as well as in negative associations for E4 and biologically younger individuals. This pattern with APOE is consistent with the known plasmalogen level decrease in AD, but the associations with delta age seem contradictory to the known decrease with aging.^53,54^ This could be explained by the U-shaped pattern of plasmalogen abundance throughout aging, with plasmalogens increasing until age 30-40 to a plateau and then decreasing with age in the elderly after around age 70.^53^ With ∼98% of the Arivale cohort being younger than 70, it is likely that plasmalogens would not yet show the decrease associated with the elderly, and higher levels would correspond to greater CA and BA.

Similarities in the constructed multi-omic atlases provide important insight as well. For example, the inter-omic association signatures of biologically older males and females are highly similar in contrast to the lack of similarity between biologically younger males and females. This implies that male and female ‘omics are more closely related in a state of perturbed health or accelerated aging than in a healthy state, which has been suggested previously.^55^ One reason for this may be the impact of sex hormones, as sex-specific testosterone and estradiol decrease with age in males and females, respectively, while luteinizing hormone and follicle stimulating hormone increase with age in both sexes.^56^ The specific inter-omic associations that were strengthened by biological oldness in both sexes seemed potentially indicative of metabolic imbalance such as insulin resistance or diabetes, which testosterone and estrogen protect against.^57,58^ For instance HbA1c was more strongly associated with central carbohydrates pyruvate and mannose.

Increased glucose was more strongly associated with increased plasma (soluble) CD163, which is a marker of inflammation and associated with the development of T2D,^59^ as well as with increased plasma HGF, which is an inflammation regulator shown to be increased in chronic disease of several organs.^60^ This trend is continued when analyzing delta age, seeing positively modified associations between glucose or HbA1c with several glycolysis and TCA metabolites including 1,5-AG, pyruvate, lactate, aconitate [cis or trans], and alpha-ketoglutarate. Of note, the interaction signatures of APOE E2 were similar to those of increased biological age. Four exactly overlapping associations were significant after multiple hypothesis correction in both male APOE E2s and biologically older males, including both HbA1c and glucose being more positively associated with phenol sulfate, a gut microbiome-produced uremic metabolite linked to albuminuria in diabetes and kidney disease.^61,62^ The other two positively modified associations were between hydroxyasparagine** and *Megasphaera*, and between FST and laureate (12:0), both also potentially highlighting an imbalance of bioenergetic pathways. Increased hydroxyasparagine abundance has been correlated to reduced kidney function,^63^ and though *Megasphaera* is among the butyrate-producing microbes generally contributing toward improved glucose homeostasis,^64–66^ *Megasphaera* abundance was found in one study to be increased in diabetics and associated with a higher fasting glucose,^67^ and in another to be increased in diabetic peripheral neuropathy and associated with higher a HOMA-IR.^68^ Increased plasma FST is associated with increased T2D risk^69^ and chronic kidney disease,^70^ though laureate (12:0) seems protective against insulin resistance, however.^71^ Similar to the signature of biological oldness and increased delta age as well, APOE ε2 exhibited positively modified associations between HbA1c and TCA metabolites fumarate and maleate, as well as between glucose and TCA metabolite aconitate [cis or trans]. Along this theme, some other significantly modified associations in APOE E2 males included the strengthened association between glucose and *Klebsiella*, a genus indicative of imbalance in the gut microbiome and known to modify the metabolome,^72^ and between triglycerides and ribitol, which disrupts central bioenergetic pathways via shifting the balance of metabolites participating in the TCA cycle, ultimately increasing glycolysis while decreasing oxidative phosphorylation.^73^

These strengthened associations in APOE E2 suggest a rewiring of bioenergetic pathways reflective of accelerated aging, such as decreased sugar catabolism, potentially by shifting from glucose to fatty acid oxidation as a source of acetyl-CoA feeding into the citric acid cycle, or an increased conversion of sugars to HbA1c in the blood. This could signify that clinically well APOE E2 individuals exhibit a signature similar to insulin resistance as compared to E3, which is supported by APOE E2’s association with increased DAGs discussed earlier. This may be indicative of E2 showing decreased preference of glucose as an energy source as compared to E3, and thus have less insulin signaling in general. Because deregulated nutrient sensing is a hallmark of aging with insulin signaling decreasing in both physiological and accelerated aging,^1^ it is unsurprising that associations suggesting insulin resistance are found in biologically older individuals. As commented upon, this connection between APOE E2 and biological oldness seems contradictory, with APOE E2 generally predicting longevity and being protective against AD, whereas insulin resistance and diabetes as suggested here are risk factors for dementia and accelerated aging.^74–76^ However, there may be an age-dependent effect of APOE, with APOE E2 imparting disadvantageous effects earlier in life while expanding longevity later. APOE E2 is associated with type III hyperlipoproteinemia^77,78^ and has been linked to increased malaria infections and severity in early childhood.^43^ On the other hand, APOE E4 shows some advantages at earlier life stages in comparison to E3 such as improved neural and cognitive development in youth and decreased infant and perinatal mortality,^43^ and was recently found to have a protective effect against obesity and T2D.^46^ This age-specific effect is an important consideration because cohort participants are relatively young and their health is representative of the US population. Therefore, E2 likely is not yet exhibiting its late-life advantages. Further, APOE E2’s potential association with insulin resistance from this analysis could suggest one of its mechanisms for supporting longevity, as a constitutive decrease in insulin signaling and insulin-like growth factor signaling would decrease the rate of cell growth and metabolism and thus reduce the rate of associated cellular damage seen in aging and AD.^1^

TwinsUK was chosen as a validation cohort because of its similarities with Arivale in being composed of community dwelling individuals and the shared usage of the Metabolon platform, enabling more direct comparison of metabolomics data. We were unable to reproduce some of our Arivale findings, including the similarities between APOE E2 and biological oldness in males, due to data limitations such as a small male sample size (maximum n = 55 for the tests of male APOE E2 and biological old males), giving low statistical power to some models and inflating the number of significant hits in biologically younger males. Some results observed in Arivale, such as biological oldness positively modifying associations between central bioenergetic metabolites and APOE E2 positively modifying the association between triglycerides and ribitol in males, trended in the same direction, however significance was lost after FDR correction. Validation results for the interaction analysis are thus uncertain. Even so, other FDR-significant interactions in TwinsUK were identified, including altered lipidomic associations in APOE E2 and in APOE E4 in females, confirming APOE exerts pressure on metabolic pathways and associations with the potential of ‘rewiring’ them to influence health status overall.

Limitations of this study include the use of cross-sectional data with no available disease or longevity outcomes to analyze. Metabolomics data was also limited to the plasma, not allowing further study on transport and localized measures such as brain metabolomics. Lifestyle factors such as diet, exercise, and medication use other than cholesterol reducing drugs were not analyzed. The available cohorts were also predominantly composed of non-Hispanic Whites (71% in Arivale, >99% in TwinsUK), which limits the generalizability of the aforementioned significant differences in APOE’s manifestation across ethnicities. Survivorship bias is another potential limitation to results for APOE E4 associations, as it well documented that older ε4 carriers represent a cognitively resilient population because many ε4 carriers die prematurely relative to ε3/ε3 individuals.^79–82^ Validation was limited by data differences between the Arivale and TwinsUK cohort, including a lower percentage and sample size of males in TwinsUK (3.6%, *n* = 61 unique individuals); lack of significant DAG species in the validation set; and lack of HbA1c measures in the validation set, having only 33 total measurements, all in females and not enough to allow all model covariates to be represented. While this study provides promising preliminary findings, future studies with greater statistical power, more diverse participants, and longitudinal data are needed to understand the universality of these results and assess the effectiveness of related interventions.

These findings substantiate APOE’s influence on bioenergetic metabolism, show agreement with current understanding and hypotheses of APOE including context dependencies such as sex differences, and suggest a mechanism for APOE-associated longevity and potentially AD pathology. Further, the results provide a preliminary atlas of inter-omic associations useful for possible interventions to offset APOE-associated risk in the prodromal stages of AD and cardiovascular disease and to extend healthspan.

## Methods

### Institutional Review Board Approval

Procedures for this study were run under the Western Institutional Review Board (WIRB) with Institutional Review Board (IRB) study number 20170658 at the Institute for Systems Biology and 1178906 at Arivale.

### Arivale Wellness Cohort and Data Collection

Research subjects in this study were voluntary, anonymous participants of the Arivale Scientific Wellness program described by Zubair et al.^26^ The program aimed to leverage the collection of dense health data from subscribers to offer personalized wellness coaching from a systems biology perspective. The collection of plasma metabolomics (Metabolon platform), plasma proteomics (Olink platform), microbiomics (16S V4 amplicon sequencing data from stool samples), and clinical chemistries data has been described thoroughly in Wilmanski et al.^83^ In this study, only individuals with whole genome sequencing were included. APOE status was determined from single nucleotide polymorphisms (SNPs) from this data, with both homozygotes for (ε2/ε2) and carriers of (ε2/ε3) the ε2 allele being defined as APOE E2; ε3/ε3 being defined as APOE E3; and both ε3/ε4 and ε4/ε4 being defined as APOE E4. ε2/ε4 individuals were excluded from analysis. The dataset contained no ε1 alleles.

### TwinsUK Cohort and Data Collection

The TwinsUK cohort was originally intended to investigate rheumatologic diseases in identical twins in the United Kingdom, and has since expanded to encompass over 15,000 volunteer identical and non-identical twins.^27^ Similar to the Arivale cohort, the voluntary participants are community dwelling, representative of the health of the population, and deeply phenotyped.

Unlike in Arivale, no coaching or intervention is performed. For this study, only the 1696 individuals with metabolomics and genotyping data were included. APOE status was determined from SNPs from Illumina assays. As in the Arivale cohort, both homozygotes for (ε2/ε2) and carriers of (ε2/ε3) the ε2 allele were defined as APOE E2; ε3/ε3 were defined as APOE E3; and both ε3/ε4 and ε4/ε4 were defined as APOE E4. ε2/ε4 individuals were recorded but excluded from analysis, and the dataset contained no ε1 alleles. Metabolomics data was obtained using the Metabolon platform, the same platform as Arivale, and has been described previously in Long et al.^28^ Demographic differences for the Arivale and TwinsUK individuals at baseline are summarized in **Table 1**

### Biological Age and Delta Age

Biological age (BA) values for the Arivale dataset were previously calculated by Earls et al.^3^. Briefly, four baseline biological age measures were computed: one from clinical labs, another from proteomics, one from metabolomics, and one combining the first three sources. Each of the models were obtained utilizing the Klemera-Doubal method, and were constructed separately for males and females. In this study, the average of the clinical lab and proteomic BA was used for metabolomic analyses, and the combined measure of BA was used for multi-omic analyses.

Chronological age (CA) at baseline was subtracted from BA to yield ‘delta age’. A delta age of over 7.5 years (about one standard deviation for both male and female, 7.8 years for female, 8.5 years for male) was treated as ‘biologically older’, and a delta age of less than −7.5 years was taken to be ‘biologically younger’. Delta age was not significantly different across APOE status (**Figure S2a)**, and sorting of APOE and delta age groups was not interdependent based on Chi2 testing (**Figure S2b**).

BA and delta age values for the TwinsUK cohort were calculated using the same modeling method used by Earls et al^3^ to calculate a metabolomics-based BA in the Arivale cohort. The 494 metabolites overlapping out of the 740 appearing in the original model were used following the same Klemera-Doubal (KD) method implemented by Earls to train another model with the TwinsUK data: BA was predicted for each sample by taking the average of ten iterations of ten-fold cross-validation, training the model separately for males and females. CA was then subtracted from BA to yield delta age. Retraining a new model independently for TwinsUK avoids errors due to batch effects. Similar to Arivale, a delta age of over 7.5 years was treated as ‘biologically older’, and a delta age of less than −7.5 years was taken to be ‘biologically younger’ for females (delta age standard deviation for females 7.8 years), however the delta age cutoff for males was set to +/- 5.0 years to better reflect the standard deviation for males (5.1 years) and smaller sample size (**Figure S3**). For TwinsUK, delta age was significantly different across APOE status (**Figure S4**).

### Differential Metabolite Abundance Analysis

For individual baseline metabolite level comparisons in Arivale, 896 winsorized metabolites from the Metabolon platform were analyzed after excluding those with more than 20% missingness. Note that isomer information for lipids such as DAGs in the Metabolon platform are not known, for example the metabolite reported as linoleoyl-arachidonoyl-glycerol (18:2/20:4) [1]* is a DAG having linoleic and arachidonic acid residues, however their position relative to the glycerol is ambiguous. Missing data was replaced via random forest imputation, which has shown to be effective for LC-MS metabolomics data.^84^ Metabolomics data was then log2 transformed. For GLMs analyzing differential metabolite abundance as reported in **Figure 2**, the model, log2(metabolite) = intercept + **APOE E2** + **APOE E4** + age + sex(Male) + BMI + cholesterol meds(self-reported) + [genetics] Principal Component (PC)1 + PC2 + e, was used for analyzing APOE, and the model, log2(metabolite) = intercept + **Biologically Young** + **Biologically Old** + age + sex(Male) + BMI + cholesterol meds(self-reported) + PC1 + PC2 + e, was used for analyzing delta age status (analyzed beta values bolded). The first two PCs used in the model were previously calculated.^83^

For TwinsUK, 752 metabolites also from the Metabolon platform remained after excluding those with more than 20% missingness. After random forest imputation, the earliest visit measurement for each individual was used after removing samples recorded as non-fasting. The following GLM model was used for analyzing APOE: log2(metabolite) = intercept + **APOE E2** + **APOE E4** + age + sex(Male) + BMI + cholesterol-reducing medications(self-reported) + batch(2-5 compared to 1) + e (analyzed beta values bolded). Delta age was not analyzed because BA models in TwinsUK were solely derived from metabolomics.

Following GLMs, an enrichment analysis of the sub-pathways annotated in the Metabolon platform was performed both on the sets of metabolites with significant (pFDR < 0.1) and pre-adjusted (*p* < 0.05) positive and negative associations for each experimental group. A standard overrepresentation test was performed for the enrichment analysis, using a hypergeometric distribution model with a survival function to calculate *p*-values for each sub-pathway for each set of associations.

### Inter-omic Interaction Analysis

For the analysis of inter-omic interactions with APOE and health, individuals were stratified by sex and then again by either APOE status or delta age status to offer direct comparisons, creating eight subsets.

For Arivale, baseline metabolomic, proteomic, and clinical chemistries data were winsorized for use in the analysis via iteratively shrinking outliers to within five standard deviations of the median. Proteomic data was from the Olink platform. Clinical chemistries were limited to only those from the Laboratory of Cell Analysis, and individuals using a different platform were dropped. Microbiome data was from DNA Genotek OMNIgene GUT collection kits sequenced by Second Genome and DNA Genotek. Baseline microbiome data was centered log-ratio transformed and filtered for rare taxa using mean and prevalence thresholds of 10 and 0.1, respectively. A total of 509,360 inter-omic combinations of analytes were tested from 876 metabolites, 274 proteins, 67 clinical draws, and 158 microbiome genera having less than 20% missing values. Those analyte pairs with significant (pFDR < 0.1) interaction results in Arivale were tested for validation in the TwinsUK cohort, given data availability. The inter-omic interactions between Glucose, LDL, HDL, Triglycerides, and Total Cholesterol from the clinical chemistries and metabolites in the ‘TCA Cycle’, ‘Glycolysis, Gluconeogenesis, and Pyruvate Metabolism’, ‘Fructose, Mannose and Galactose Metabolism’, ‘Pentose Metabolism’, ‘Oxidative Phosphorylation’, ‘Phospholipid Metabolism’, ‘Sphingolipid Metabolism’, ‘Leucine, Isoleucine and Valine Metabolism’, or ‘Diacylglycerol’ subpathways (Metabolon labeling) were additionally analyzed in the validation cohort. TwinsUK data preprocessing followed the same method as in Arivale, with a 20% missingness threshold and iterative winsorization of outliers to within 5 standard deviations of the median. All samples indicating non-fasting were dropped, and the TwinsUK data from the earliest visit containing values for both clinical test and metabolite for each individual were used in the analysis. Intra-omic combinations were not tested in either cohort.

GLMs were performed for each subset with the following model for Arivale: analyte1 = intercept + analyte2 + X + **analyte2*X** + age + season(reference = Fall) + BMI + cholesterol meds(self-reported) + [genetics] Principal Component (PC)1 + PC2 + e, where X is the experimental group analyzed (ie: APOE E2 or E4, or Biologically Young or Old statuses) and ‘analyte2*X’ represents the interaction term between the second analyte and the experimental group, bolded here to indicate it is the beta value analyzed. The first two PCs used in the model were previously calculated.^83^ For TwinsUK, the model was: clinical test = intercept + metabolite + X + **metabolite*X** + age + BMI + cholesterol meds(user) + e. Each experimental group was analyzed separately and stratified by sex to isolate and narrow focus on the variable of interest. An additional set of models was tested in each cohort as well, with sex as a covariate instead of a stratified variable, and the ‘experimental groups’ being ε2 or ε4 allele dosage (with possible values being 0, 1, or 2) or delta age value in days. The relatively small number of analyte pairs in the GLMs failing with a ‘NaN, inf or invalid value detected in weights, estimation infeasible’ error, were noted but ignored.

All GLM models in statistical analysis assumed a gaussian distribution with an identity link and were set at 2000 maximum iterations. For the inter-omic interaction analysis, if the first analyte exhibited a skew of greater magnitude than 1.5, a gamma distribution was used with a log link instead, with values of zero being replaced with half the minimum non-zero value. FDR significance was determined by adjusting *p*-values corresponding to APOE and health statuses by the Benjamini-Hochberg method with the FDR set to 5%.^85^ GLMs were performed using the statsmodels package version 0.13.0 in Python version 3.9.7.

## Acknowledgements

The authors would like to thank Max Robinson, and other lab group members and collaborators that shared thoughtful comments and the participants of the Arivale Wellness Program and TwinsUK who consented to let their deidentified data be used for research purposes. This work was funded by a generous gift from K. C. Ellison (to K.W. and T.W.); Japan Science and Technology Agency (JST) PRESTO Program (JPMJPR238A to K.W.); the Global Grants for Gut Health from Nature Portfolio and Yakult (to S.M.G.); National Institutes of Health (NIH) grants no. U19AG023122 & Translational Opportunity Fund (N.R, O.F, P.S), R01AG061844 (P.S), and 1K12TR004384 (A.A.K); and USDA Agricultural Research Service under Cooperative Agreement No. 58-8050-3-003 (A.A.K). TwinsUK is funded by the Wellcome Trust, Medical Research Council, Versus Arthritis, European Union Horizon 2020, Chronic Disease Research Foundation (CDRF), Wellcome Leap Dynamic Resilience Programme (co-funded by Temasek Trust), Zoe Ltd, the National Institute for Health and Care Research (NIHR) Clinical Research Network (CRN) and Biomedical Research Centre based at Guy’s and St Thomas’ NHS Foundation Trust in partnership with King’s College London. We thank Allison Kudla for help in graphic design.

## Author Contributions

D.E, C.F, P.B and N.R, conceptualized the study. D.E, K.W, T.W, and G.G performed data analysis and figure generation. M.S.L, K.W., J.J.H, O.F, P.S, N.D.P, L.H, S.J.E, L.P, J.C.L, S.M.G, D.E, C.F, P.B, A.A.K, and N.R assisted in results interpretation, A.T.M managed the logistics of data collection and integration, L.P provided code used in model building. D.E and N.R were the primary authors of the paper, with contributions from all other authors. All authors read and approved the final paper.

## Conflicts of Interest

The authors declare no conflict of interest.

## Supplementary Files

**Supplementary File 1** contains the results data for the discovery cohort (Arivale) individual metabolite abundance analysis and corresponding enrichment analysis.

**Supplementary File 2** contains the results data having pFDR < 0.1 for the discovery cohort (Arivale) inter-omic interaction analysis.

**Supplementary File 3** contains the results data for the validation cohort (TwinsUK) individual metabolite abundance analysis and corresponding enrichment analysis.

**Supplementary File 4** contains the results data for the validation cohort (TwinsUK) inter-omic interaction analysis.

**Figure S1:**
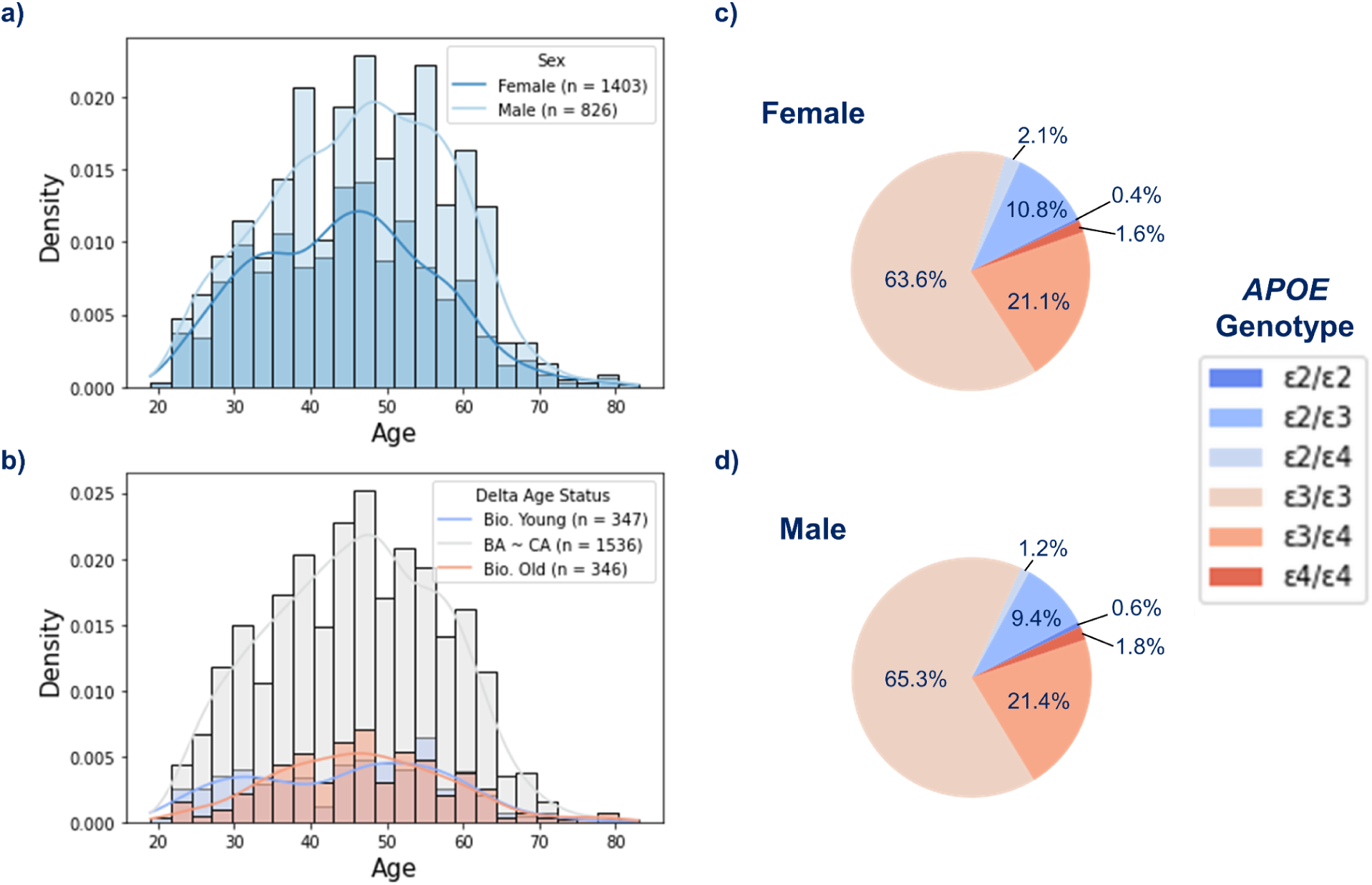
The Arivale cohort contains a range of community dwelling individuals spread across ages, delta age statuses, and APOE genotypes. (**a**, **b**) Density histograms of baseline chronological ages in the Arivale cohort stratified by sex (**a**) and delta age status, with biologically young and old defined as having a biological age 7.5 years younger or older than chronological age, respectively (**b**). The lines indicate the kernel density estimates. **c**, **d** Pie charts displaying APOE genotype frequencies in the female (**c**) and male (**d**) Arivale participants. Presented is the baseline data used in interaction analyses.

**Figure S2:**
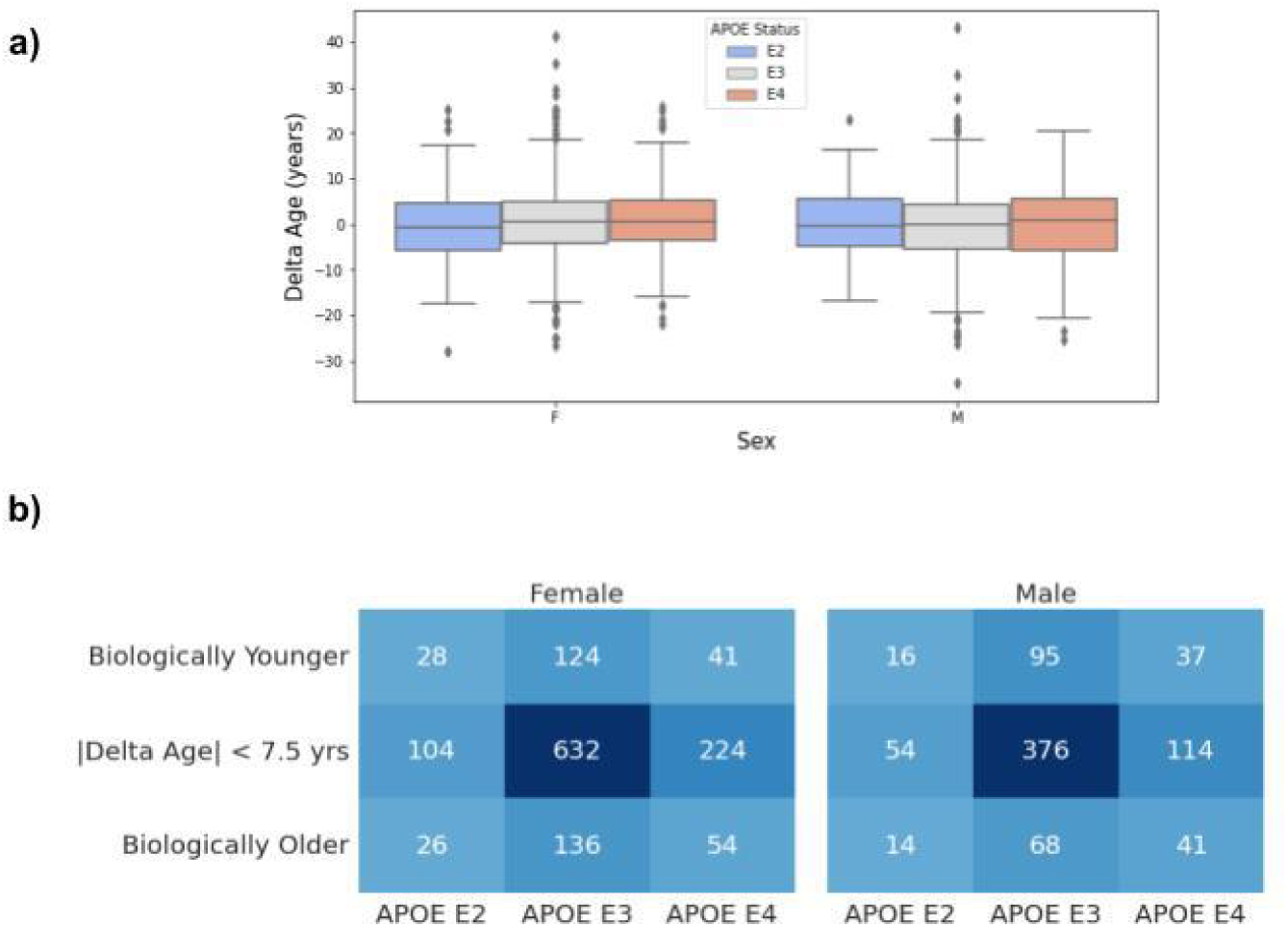
Delta age and delta age-based stratifications are not significantly different across APOE statuses for either males or females in Arivale. (**a**) Box plot of delta age across APOE statuses. Pairwise Mann–Whitney U tests between APOE statuses within male and female showed non-significant *p*-values (smallest *p*-value = 0.062 between female E2 and E4). *n* = 158 (Female E2), *n* = 892 (Female E3), *n* = 319 (Female E4), *n* = 84 (Male E2), *n* = 539 (Male E3), *n* = 192 (Male E4). (**b**) Counts of individuals in APOE and delta age categories, stratified by sex. The chi-squared tests yielded *p* = 0.59 for females and *p* = 0.041 for males.

**Figure S3:**
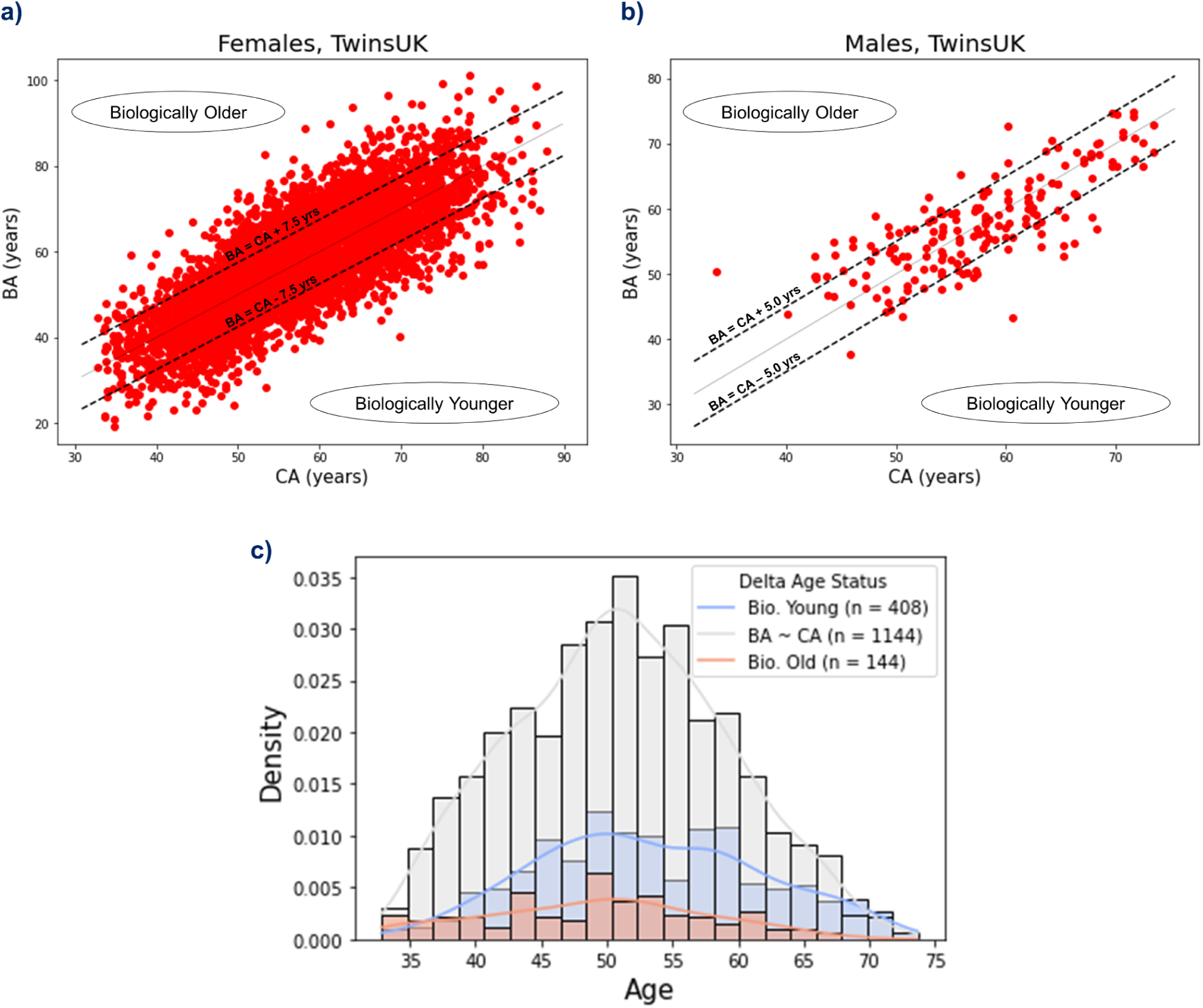
Metabolomic BA was predicted by fitting a model to TwinsUK data. (**a**, **b**) The scatterplot of BA and CA for female (*n* = 1,635 individuals, Pearson’s r = 0.778) (**a**) and male (*n* = 61, r = 0.776) (**b**) TwinsUK participants. The solid line indicates BA = CA, and the dotted lines indicate cutoffs for defining the Biologically Younger and Older groups. See Methods for model details. (**c**) A density histogram of baseline chronological ages in the TwinsUK cohort stratified by delta age status. The lines indicate the kernel density estimates.

**Figure S4:**
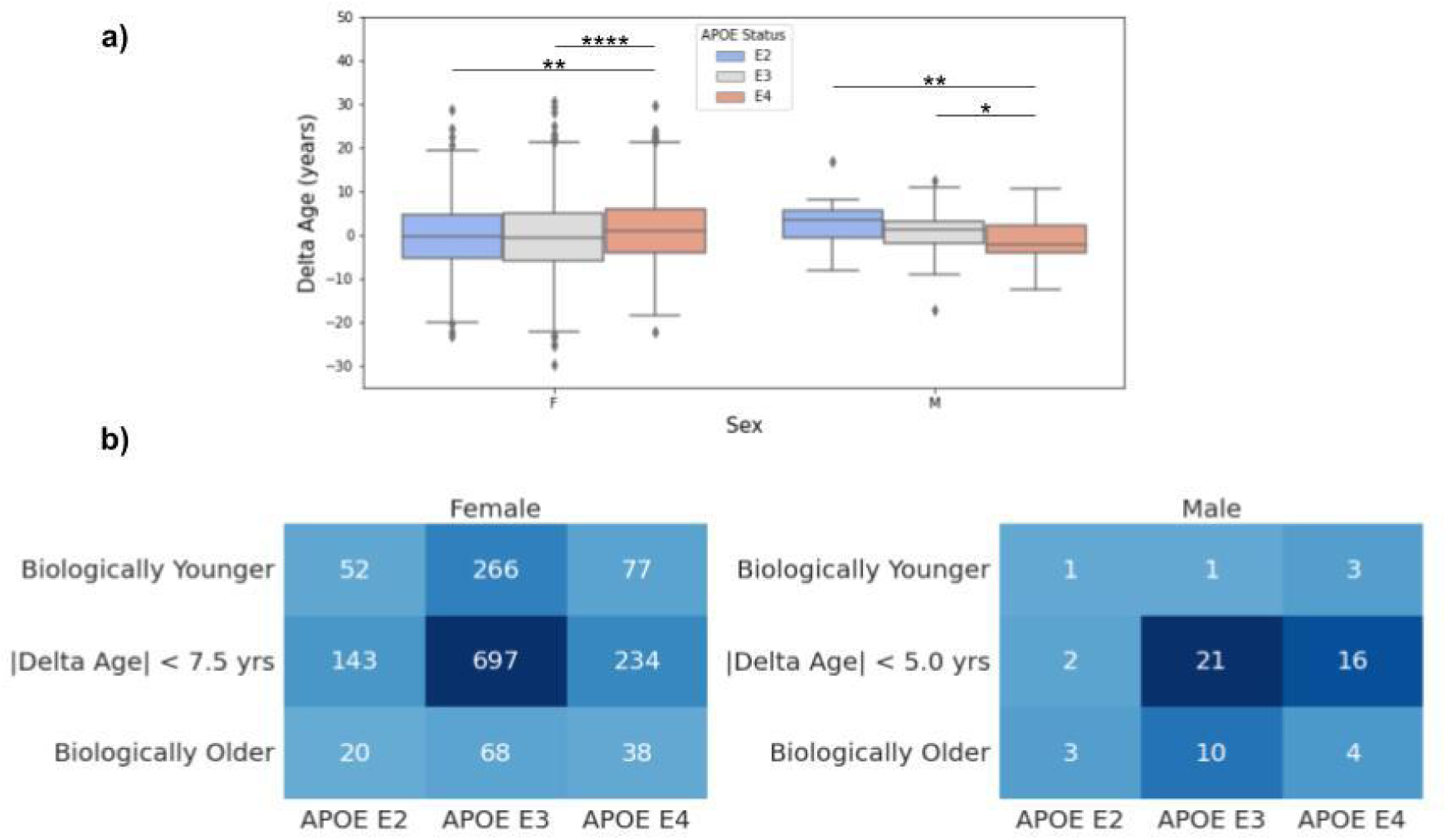
Delta age is significantly different across APOE statuses in the TwinsUK cohort. (**a**) Box plot of delta age across APOE statuses for all TwinsUK participants, including longitudinal. \**p*<0.05, \*\**p*<0.01, \*\*\**p*<0.001, \*\*\*\**p*<0.00001 based on pairwise Mann-Whitney U-tests. *n* = 641 (Female E2), *n* = 3050 (Female E3), *n* = 1043 (Female E4), *n* = 18 (Male E2), *n* = 95 (Male E3), *n* = 69 (Male E4). (**b**) Counts of individuals in APOE and delta age categories at baseline visit, stratified by sex. The chi-squared tests yielded *p* = 0.26 for females and *p* = 0.08 for males.

**Figure S5:**
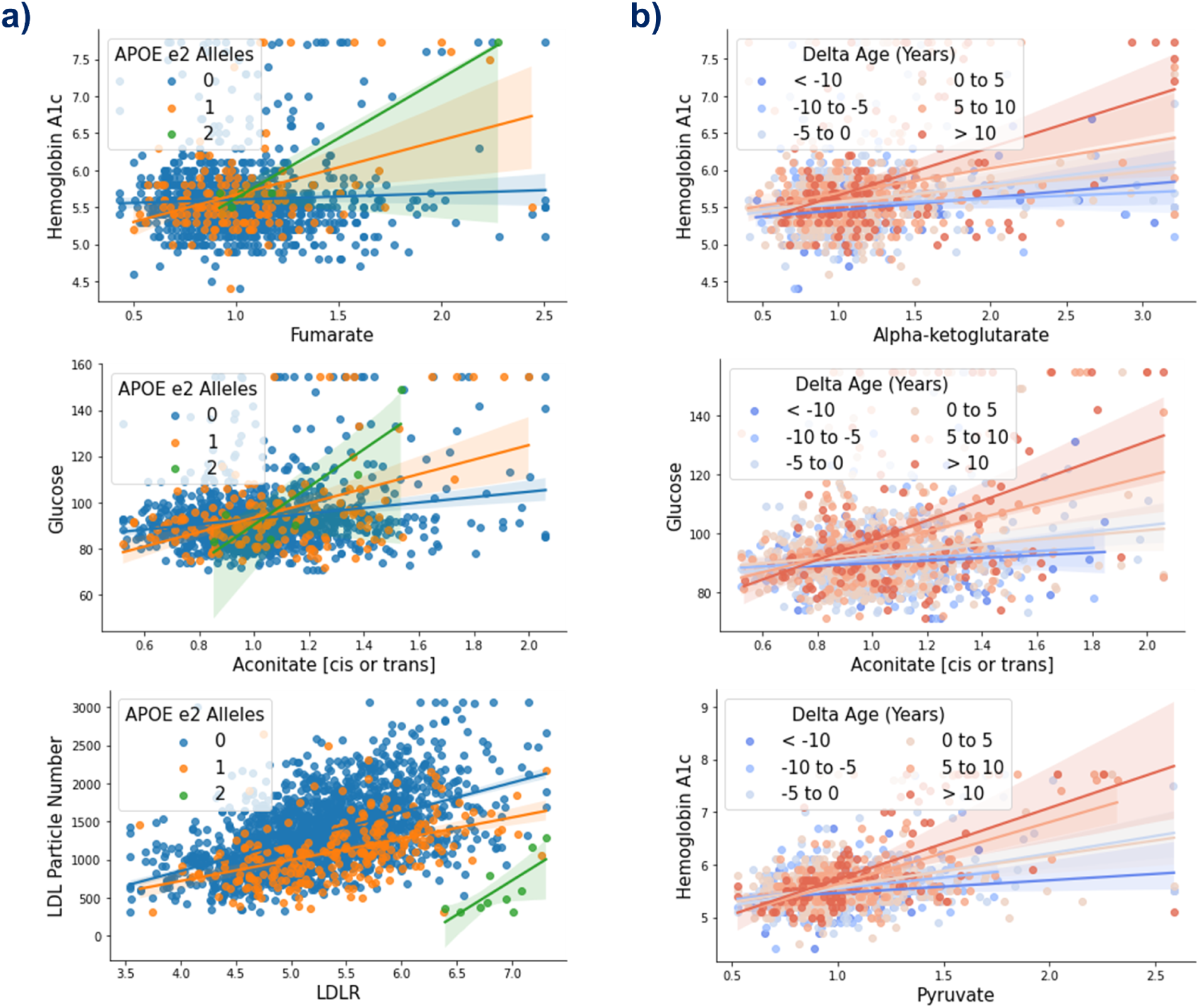
Inter-omic associations are modified by ε2 allele dosage and continuous delta age. Scatter plots of inter-omic analyte pairs with associations significantly modified by APOE ε2 allele dosage (**a**) and by delta age (**b**). Line indicates simple linear regression, with shading indicating the 95% confidence interval.

**Figure S6:**
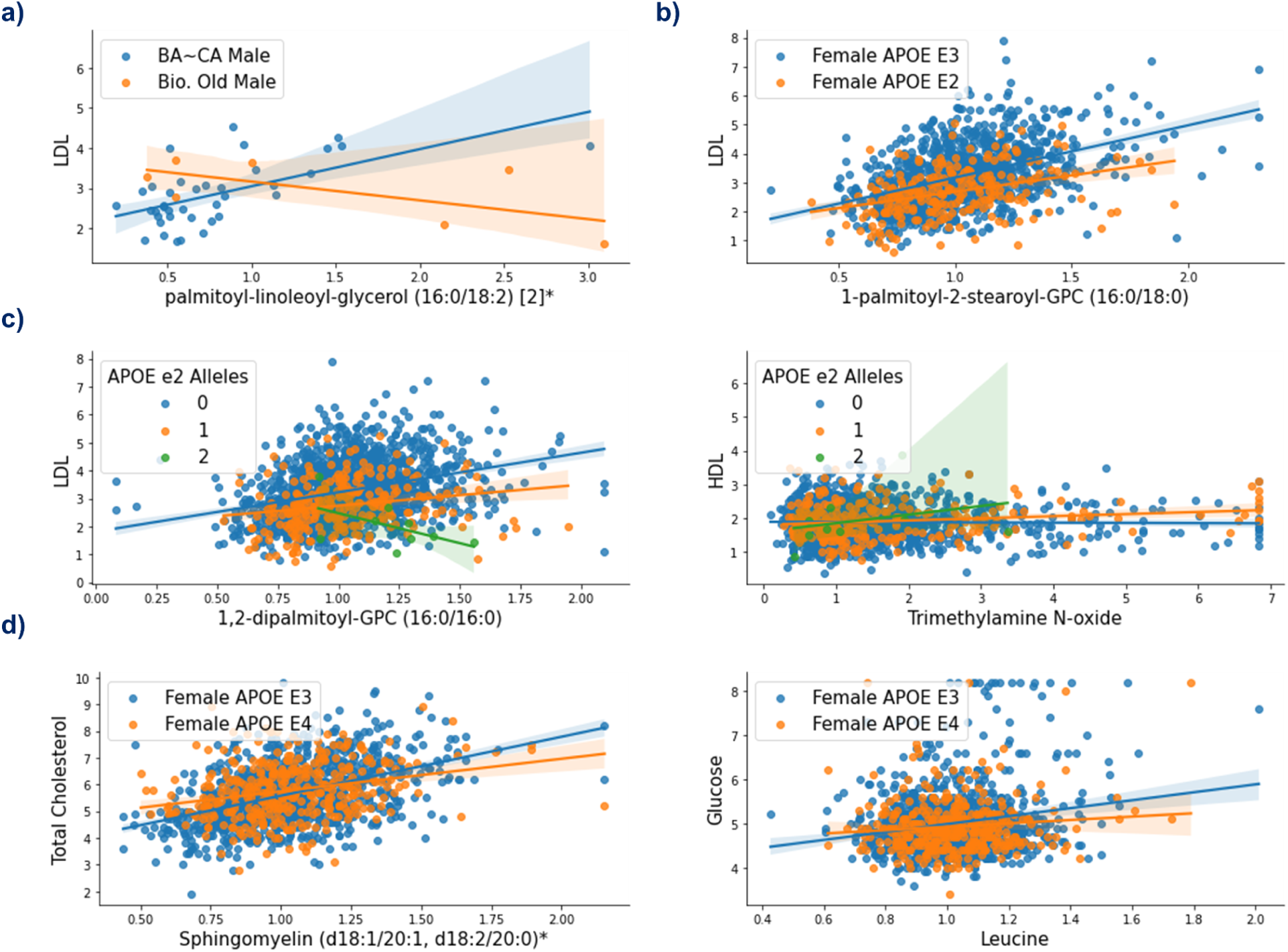
Inter-omic associations involving lipids are modified by APOE and delta age statuses in TwinsUK validation cohort. Scatter plots of inter-omic analyte pairs with associations significantly modified by biological oldness in males (**a**), APOE E2 in females (**b**), the APOE ε2 allele (**c**), APOE E4 in females (**d**) in the TwinsUK cohort. Line indicates simple linear regression, with shading indicating the 95% confidence interval.

**Supplementary Table 1:**
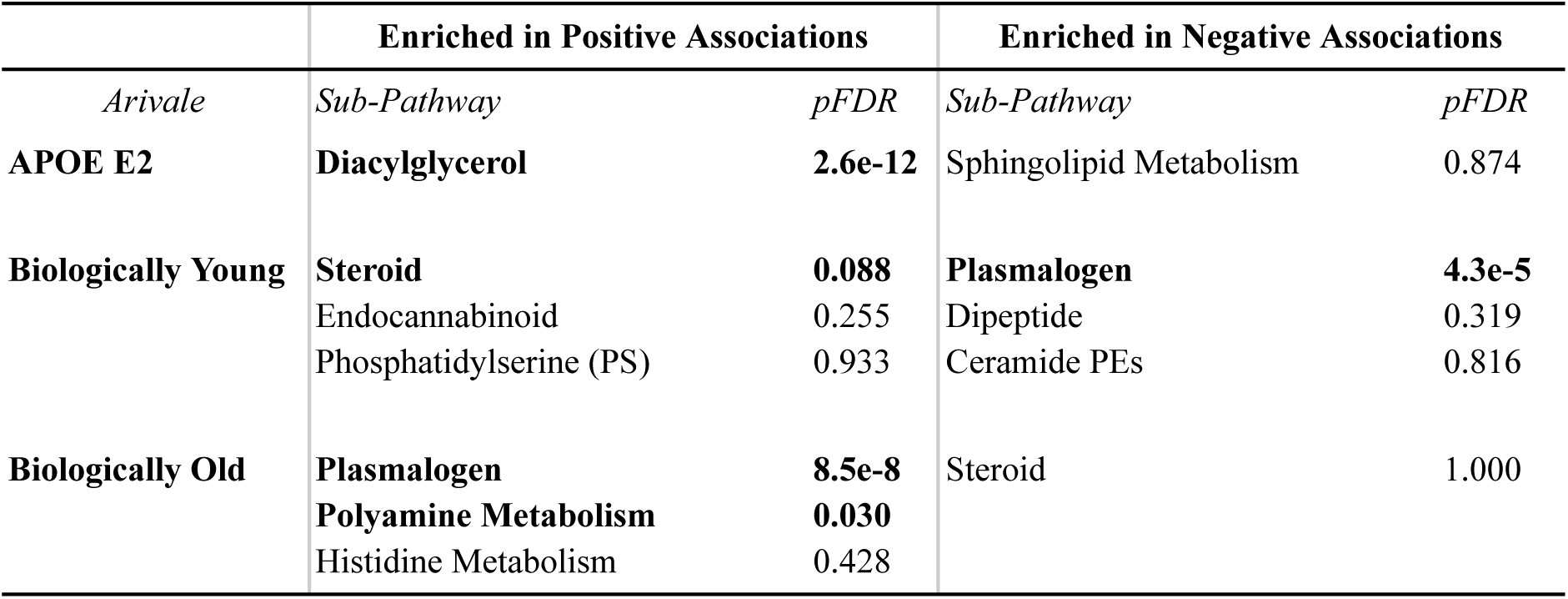
Enriched metabolic sub-pathways in the metabolites significantly associated with APOE or delta age statuses. Presented are the metabolite sub-pathways, as categorized by the Metabolon platform, enriched with *p* < 0.05 in the metabolites that exhibited significantly positive or negative associations with APOE or delta age statuses after FDR correction (pFDR < 0.1). Bolding denotes pFDR < 0.1 (Benjamini–Hochberg method) of the enrichment analysis.

**Supplementary Table 2:**
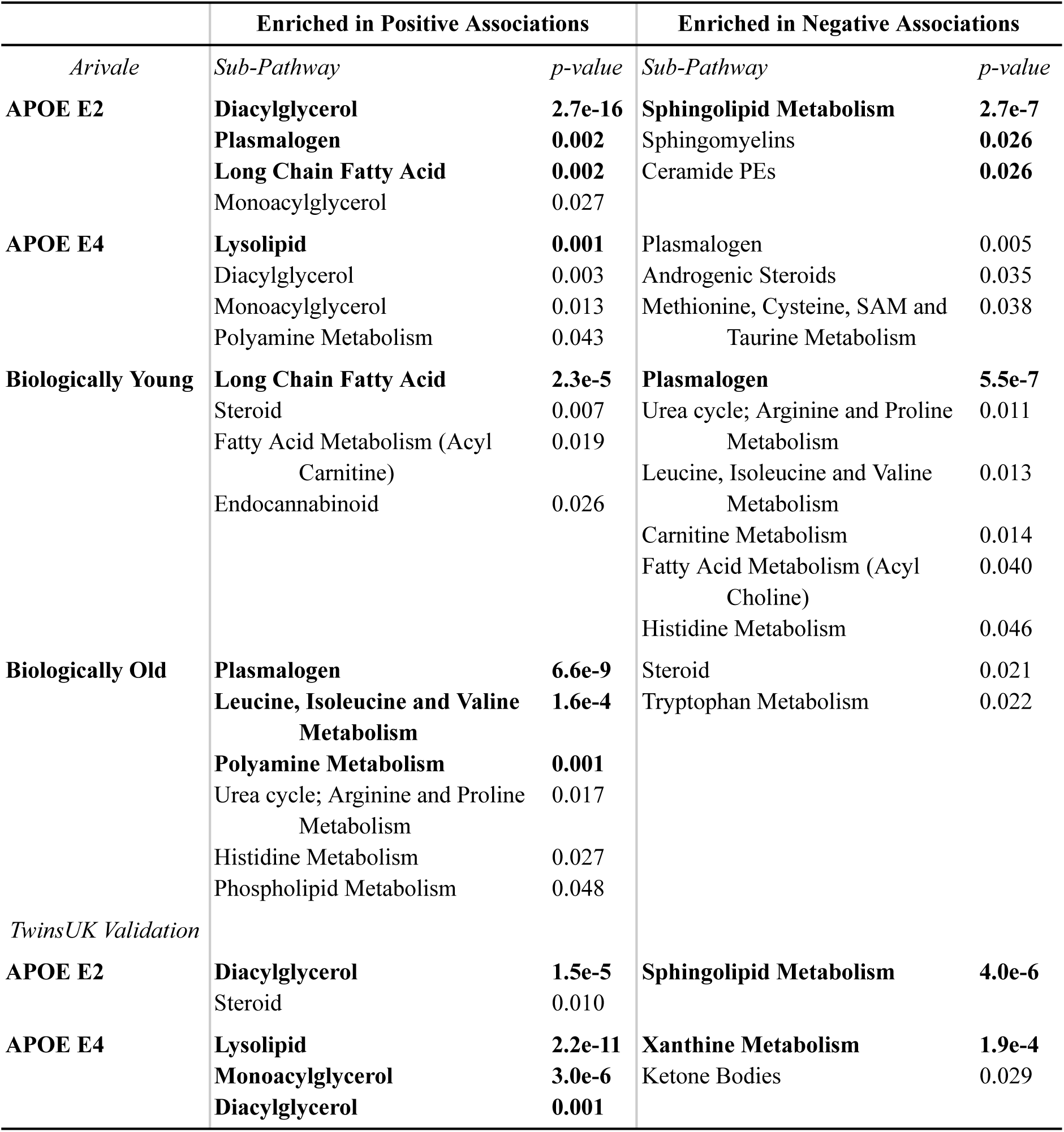
Enriched metabolic sub-pathways in the metabolites associated pre-adjustment with APOE or delta age statuses. Presented are the metabolite sub-pathways, as categorized by the Metabolon platform, enriched with *p* < 0.05 in the metabolites that exhibited positive or negative associations with APOE or delta age statuses with pre-adjusted *p* < 0.05. Bolding denotes pFDR < 0.1 (Benjamini–Hochberg method) of the enrichment analysis.

**Supplementary Table 3:**
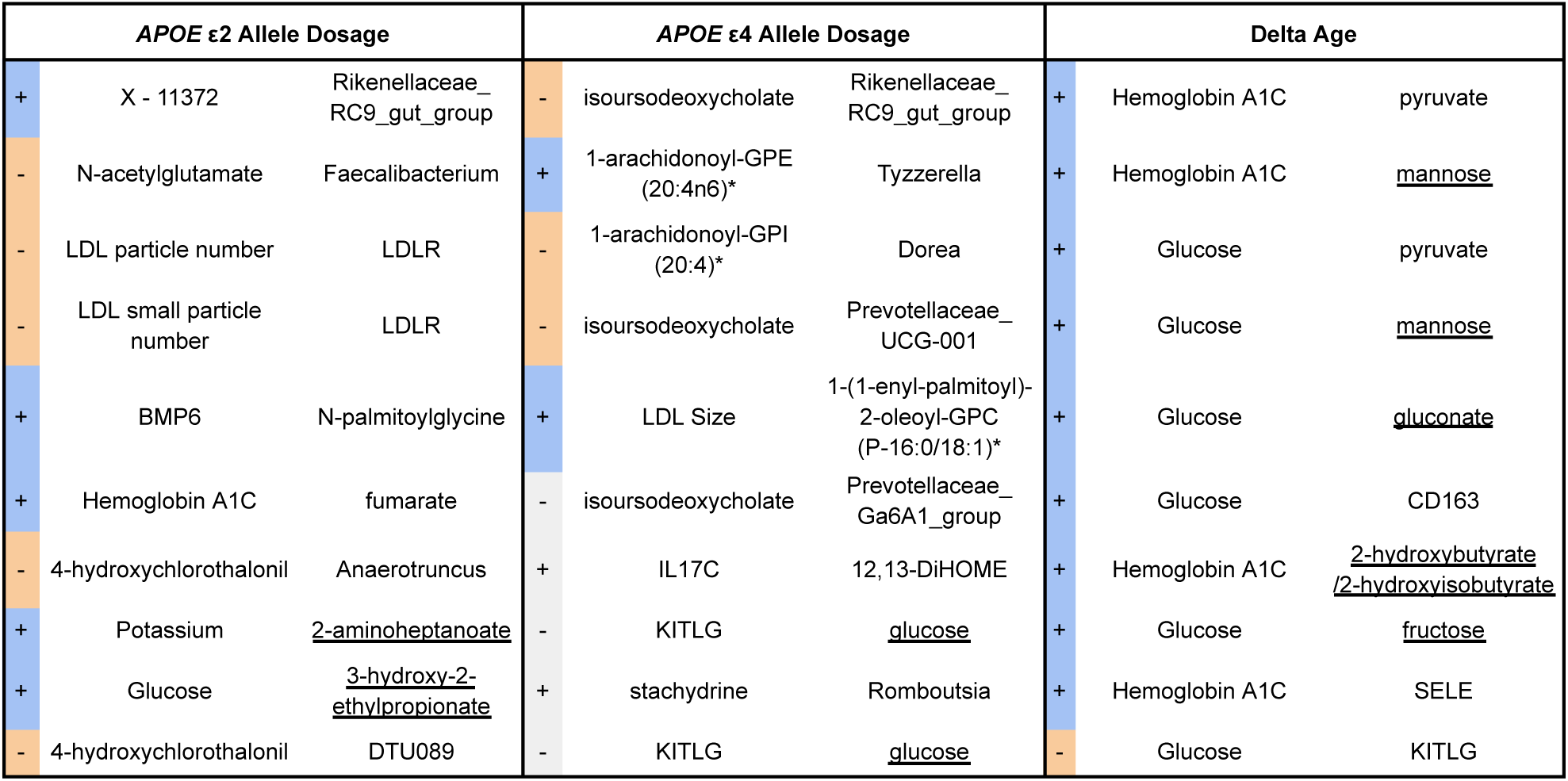
Top ten inter-omic analyte pair associations modified by APOE allele dosage and delta age in Arivale. For each set of models, the ten analyte pairs with the lowest *p*-values for the interaction term representing the modification of APOE allele dosage or delta age on the association between the two analytes are tabulated. Blue and orange highlighting indicate positive and negative interaction terms, respectively, while gray indicates pFDR > 0.1 (**Supplementary File 2** for full data). Underlining indicates a metabolite associated with the experimental group in the analysis of differential metabolite abundance (with pre-adjusted *p* < 0.05).

## Notes

### Competing Interest Statement

The authors have declared no competing interest.

### Summary of Updates

ORCID numbers have been added for some co-authors. Manuscript PDF is unchanged.

